# Systemic Dosing of Virus-derived Serpin Improves Survival and Immunothrombotic Damage in Murine Colitis

**DOI:** 10.1101/2024.01.08.574715

**Authors:** Liqiang Zhang, Julie Turk, Henna Monder, Cheyanne Woodrow, Laurel Spaccarelli, Aman Garg, Jessika Schlievert, Nora Elmadbouly, Aashika Dupati, Emily Aliskevich, Rohan Saju, Jacqueline Kilbourne, Kenneth Lowe, Mostafa Hamada, Aubrey Pinteric, Isabela R. Zanetti, Ritvik Srivant Satyanarayanan, Junior Enow, Esther Florsheim, Masmudur Rahman, James Irving, Grant McFadden, Wei Kong, Alexandra R. Lucas

## Abstract

Inflammatory bowel disease (IBD) is potentially life-threatening, with risk of bleeding, clotting, infection, sepsis, cancer and toxic megacolon. Systemic and local immune and coagulation dysfunction increase IBD severity. Current treatments are partially effective, but there is no definitive cure. *Ser*ine *p*rotease cascades activate thrombotic, thrombolytic and complement pathways and are regulated by *in*hibitors, *serpins*. Viruses encode proteins evolved from endogenous central regulatory pathways. A purified secreted Myxomavirus-derived serpin, Serp-1, dosed as a systemic anti-inflammatory drug, has proven efficacy in vascular and inflammatory disorders. PEGylated Serp-1 protein (PEGSerp-1) has improved efficacy in lupus and SARS-CoV-2 models. We examined PEGSerp-1 treatment in a mouse Dextran Sodium Sulfate (DSS) colitis model. Prophylactic PEGSerp-1 significantly improved survival in acute severe 4-5% DSS colitis, reducing inflammation and crypt damage in acute 4-5% DSS induced colitis and when dosed as a chronic delayed treatment for recurrent 2% DSS colitis. PEGSerp-1 reduced iNOS^+^ M1 macrophage invasion, damage to crypt architecture and vascular inflammation with decreased uPAR, fXa, fibrinogen and complement activation. This work supports PEGSerp-1 as a tissue targeting serpin therapeutic.

## Introduction

Vascular inflammation in the gut and throughout the body is closely linked to exacerbation and complications in inflammatory bowel disease (IBD) (1–9), whose global annual incidence has been estimated to be around 7 million (https://doi.org/10.1016/S2468-1253(19)30333-4). Of the two primary manifestations of this condition, Crohn’s disease (CD) is characterized by transmural inflammation that can affect any region of the gastrointestinal tract, whereas the inflammation in ulcerative colitis (UC) is more superficial and limited to the rectum and colon. In severe IBD there is an elevated risk for both clotting and bleeding, with ongoing immune-mediated damage in the gut and in the vasculature, with relapsing chronic inflammation of the gastrointestinal tract driven by neutrophils, macrophages and T cells (1–16).

The pathogenesis of IBD is not fully understood, but it is accepted that genetic susceptibility, microvascular inflammation and ischemia, and altered microbiome composition can induce or exacerbate IBD, resulting in a dysregulated immune response (1–16). Increased levels of urokinase-type plasminogen activator (uPA) and uPA receptor (uPAR), as well as fibrinogen are also detected in the colon and in the circulation in IBD (7–10), underscoring the dysregulated inflammatory and coagulopathic responses. Vascular inflammation and micro-thrombosis with reduced blood flow and ischemia are associated with IBD severity (1–3). Active IBD is also complicated by accelerated atherosclerotic cardiovascular disease and increased risk for thrombotic arterial occlusions (heart attacks) and venous thrombosis (DVT). Immune and coagulation pathways throughout the vascular system are driven by serine protease enzyme cascades in the thrombotic and thrombolytic (coagulation) (7–10) and complement pathways (11–16). *Ser*ine *p*rotease *in*hibitors, termed *serpins*, are important endogenous regulators of these protease pathways (17). Serpins are unique in that they act as suicide substrates, forming an irreversible covalent 1:1 complex with a target protease (17,18). Interdependent co-activation of the thrombolytic and thrombotic pathways together with activation of complement pathways and inflammatory responses leads to ongoing vascular and colon damage, increasing clotting and/or bleeding (8–10,15 19-21). These dysfunctional immune-coagulopathic responses can create repetitive cycles of disease. Vascular inflammation and thrombosis are thus linked to IBD complications with increased risk of thromboembolic events. Prior studies have identified altered expression of thrombolytic proteases, urokinase- and tissue-type plasminogen activators (uPA and tPA), the soluble uPA receptor (suPAR) and the clotting protease fibrin in areas of active colitis and suPAR in the blood, implicating both the thrombolytic and thrombotic arms of the coagulation cascades in disease progression (10,21). The uPA and tPA proteases are part of the inflammatory cell response, activating plasmin which in turn activates matrix metalloproteinases (MMPs)(22–25). MMPs break down connective tissue and allow immune cells to invade the vascular wall and gut tissues, increasing inflammation. Similar bidirectional activation of coagulation and immune responses is seen in severe unstable coronary diseases, such as unstable atherosclerotic plaque leading to arterial thrombosis and vessel occlusion, as in myocardial infarctions.

Altered expression of mammalian serpins, specifically plasminogen activator inhibitor-1 (PAI-1, SERPINE1), vaspin (SERPINA12), pigment epithelial derived factor (PEDF, SERPINF1), C1-inhibitor (C1INH, SERPING1) alpha-1-antitrypsin (A1AT, SERPINA1), and antithrombin III (ATIII, SERPINC1), are reported in IBD and in vascular disease (15,26–29), suggesting a role in their pathogenesis.

Therapeutics that target enzyme activity in thrombosis (heparin) as well as thrombolysis (uPA inhibitors) have both been assessed, with partial success in treating IBD (10,22). In practice, aspirin (5-aminosalicylic acid) derivatives are used as a first line treatment for IBD but are often unable to control the disease, necessitating the use of corticosteroids. Corticosteroids are associated with multiple complications when used as a long-term therapy (2–4,22), including steroid-induced diabetes, infection, vascular disease, ischemia and osteonecrosis with attendant morbidity and mortality. Other medications include azathioprine and methotrexate, as well as newer biologics that include anti–TNF-α antibodies, anti-integrin antibodies including a biologic agent against the p40 subunit of interleukin-12/23, a Janus kinase (JAK) inhibitor, and a sphingosine-1-phosphate (S1P)–receptor modulator (19). However, there remains no cure and IBD requires lifelong management, with 15%-30% of UC patients and 70-80% of CD patients requiring surgical removal of the affected region of the bowel at some time to treat complications or achieve remission (1–3,22).

Mammalian serpins regulate the thrombotic and thrombolytic coagulation cascades, as well as the complement pathways, representing up to 10% of circulating blood proteins. These function as inhibitors, targeting sites of protease activation (17,18). Viruses encode proteins co-opted and evolved to sustain cycles of infection and propagation; in some cases direct or incidental endogenous targets of these proteins mediate roles in the pathogenesis of endogenous or autogenic conditions. Poxviruses and herpesviruses are large DNA viruses that have evolved immunomodulatory proteins designed to regulate the host immune response to viral replication (17,23–25). Myxomavirus (MyxV) is a poxvirus which infects rabbits, but which is non-pathogenic to humans. Serp-1 is a secreted 55kDa glycosylated MyxV-derived serpin that binds and inhibits tPA, uPA, and plasmin in the thrombolytic cascade, and factor Xa (fXa) and thrombin in the thrombotic cascade (23–25,30–34). Additionally, Serp-1 targets several complement proteases including C1s (25,33), and more recently demonstrated to bind a series of complement proteases with reduced complement membrane attack complex (MAC, C5b/9) detection after Serp-1 treatment in animal models of inflammatory disease (33,34). Outside of the context of a viral infection, these modulatory effects on inflammation represent clinically desirable properties that give this molecule therapeutic potential. Accordingly, Serp-1 as well as a modified polyethylene glycol-conjugated (PEGylated) form (PEGSerp-1) have demonstrated efficacy in a wide array of animal models of disease (30–34). Serp-1 treatment has also proved safe and effective in a dose escalating, randomized, and blinded Phase IIA clinical trial in patients with unstable angina and coronary stent implants performed at 7 sites in the US and Canada (17,35). The modified PEGSerp-1 form shows improved half-life and efficacy, reducing inflammation and lung consolidation in a pristane-induced lupus lung hemorrhage (DAH) model (26) and in a MA30 SARS-CoV-2 infected mouse model (33,34).

Here we report a series of studies examining PEGSerp-1 treatment in mouse models of acute and chronic dextran sodium sulfate (DSS)-induced ulcerative colitis (36,37) with both prophylactic as well as delayed treatments. Clinical score, survival, and immune and coagulopathic responses were evaluated in each study. The results reported here highlight the potential for PEGSerp-1 as a treatment for IBD.

## Results

### Acute prophylactic treatment with PEGSerp-1 improves survival and colon damage in severe 4 and 5% DSS colitis

PEGSerp-1 treatment given at the time of inducing colitis, was first assessed as an acute preventative treatment in two initial studies of 4% and 5% DSS induced colitis, referred to as acute prophylaxis. PEGSerp-1 was given systemically as daily intraperitoneal (IP) injections starting on the same day that DSS was introduced in the drinking water (Figure 1A). One group of mice were challenged with 5% DSS for 6 days and a second group with 4% DSS for 5 days (36,37). With Saline vehicle treatments in the 4% and 5% DSS colitis groups, there was high mortality; 100% by day 6 with 5% DSS and 40% by day 10 for 4% DSS. PEGSerp-1 markedly improved survival when given on the first day of 5% DSS induced colitis reducing mortality to 20% (Figure 2A, *p* < 0.0326 Kapplan Meier survival). All 5 saline treated mice given 5% DSS died by day 6 while 4/5 PEGSerp-1 treated mice survived until the end of the study (day 14). There was a non-significant trend toward improved survival with PEGSerp-1 treatment in the 4% DSS group (Figure 2B, Mantel-Cox, *p* = 0.4132). Combined survival for the 4 and 5% DSS acute prophylaxis groups was also assessed and again demonstrated significantly improved survival with early prophylactic PEGSerp-1 treatment (Figure 2C, *p* < 0.0225) indicating improved survival when combining survival data for both 4 and 5% DSS.

**Figure 1.**
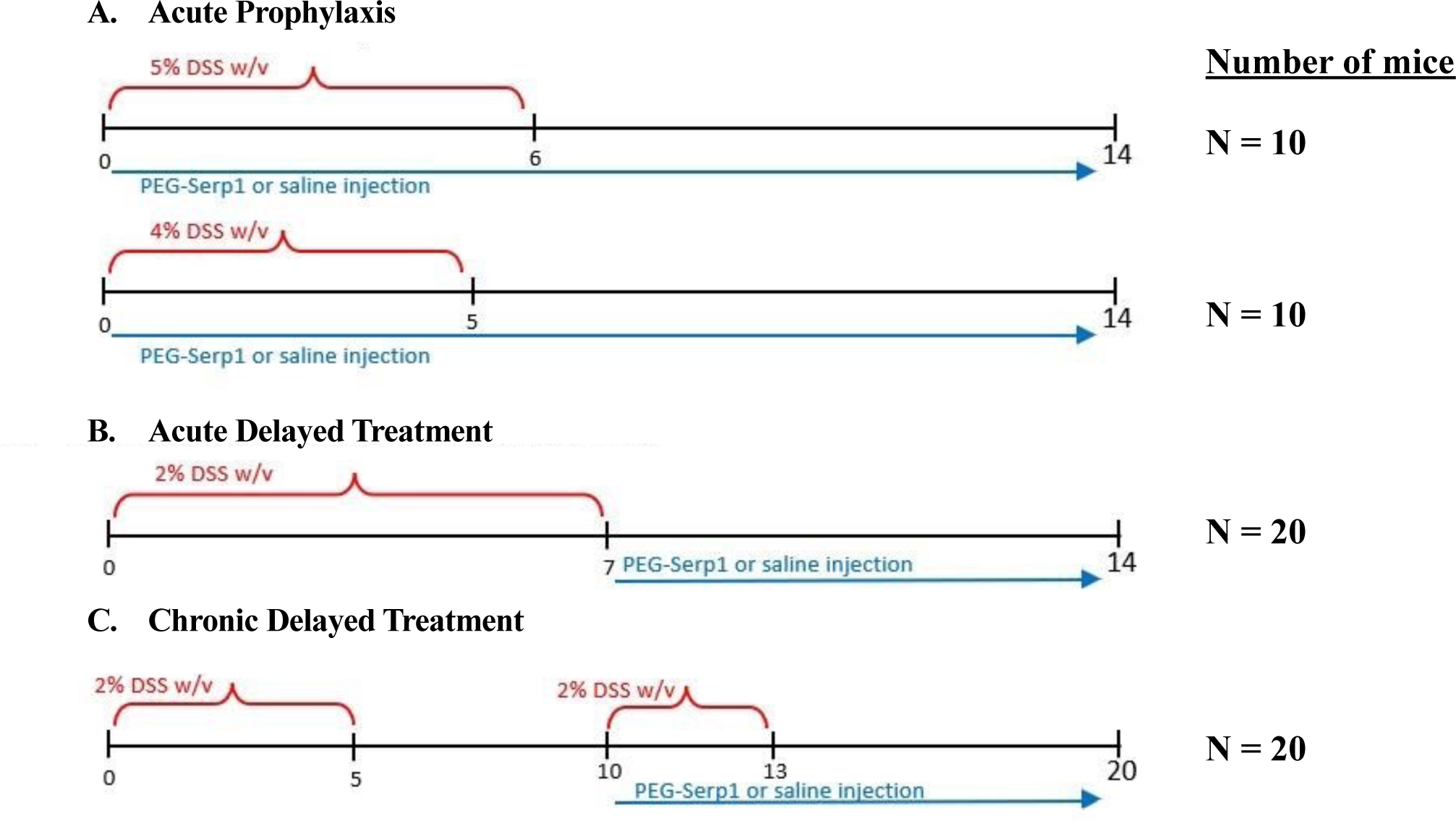
Flow chart for DSS models: **A** - Acute prophylaxis treatment models with immediate early PEGSerp-1 treatment starting on day 0 when DSS is introduced into the drinking water. **B** - 2 Acute delayed treatment starting 7 days after introducing 2% DSS into the drinking water. **C** - Chromic delayed PEGSerp-1 treatment starting on day 10 when 2% DSS is reintroduced after an initial 5days DSS and 5 days rest.

**Figure 2.**
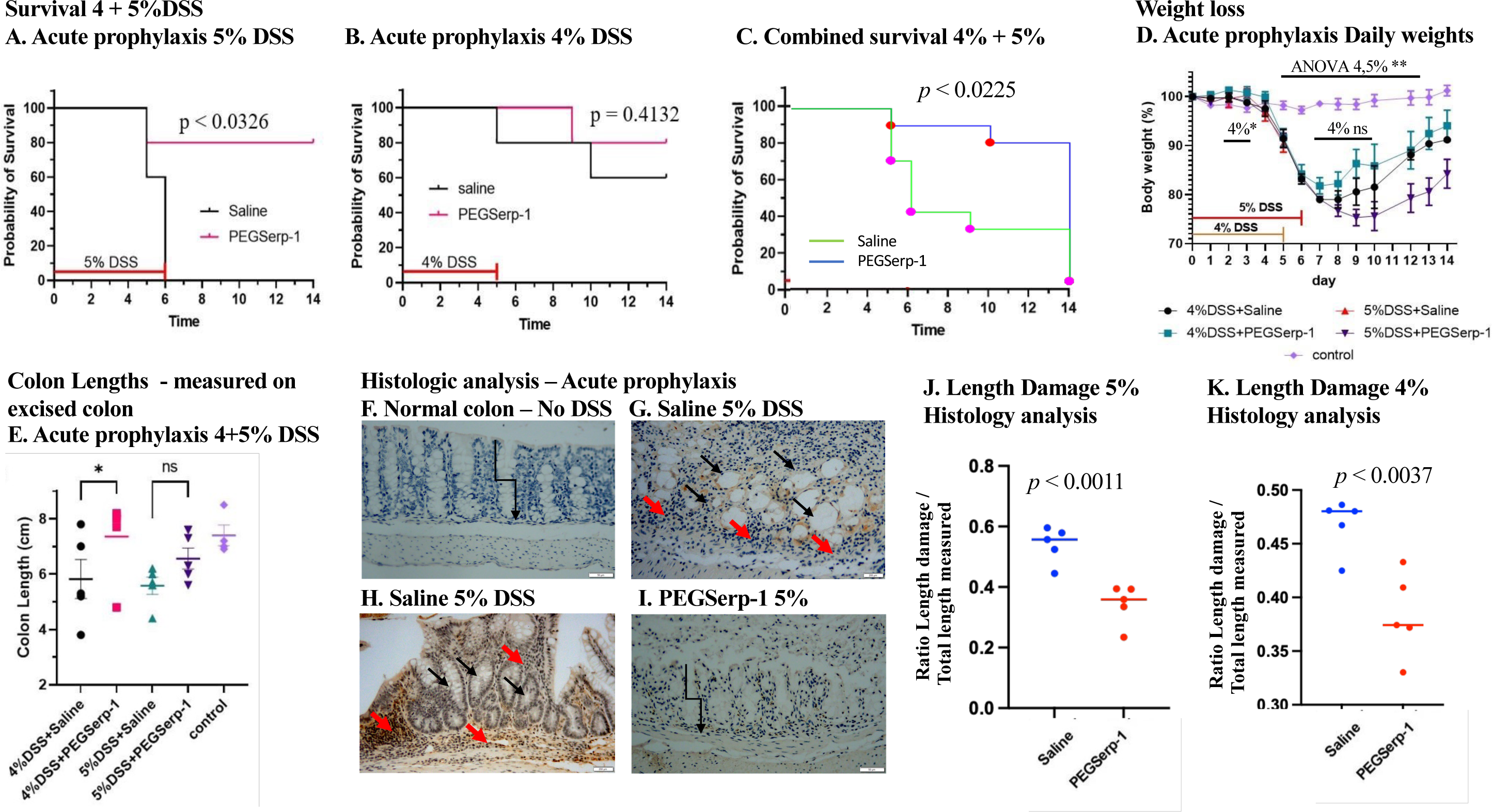
Acute prophylactic treatment with PEGSerp-1 initiated on the first day of introduction of 5% DSS improves survival (Kaplan-Meier survival analysis) (**A**, *p* < 0.0326), with a trend toward improving survival with introduction of 4% DSS in the drinking water (**B**, *p* = 0.4132). Combined survival analysis for the 4% and 5% DSS acute prophylaxis models demonstrated improved survival (**C**, *p* < 0.0225). Weight loss was significantly greater in all 4% and 5% DSS treatment groups when compared to the normal controls on days 5-12 (**D**, *p* < 0.01). Weight loss is modestly improved in PEGSerp-1 treated mice with 4% DSS at days 2-4 with no significant changes after day 4 (**D**). Due to high early mortality with 5% DSS, weight loss was only compared in the 5% DSS acute prophylaxis model from days 0 to 5 with no significant change detected (**D**). Colon length measured immediately after euthanasia and colon excision indicated increased colon length and reduced damage in 4% and 5% DSS colitis with PEGSerp-1 treatment (**E**, *p* < 0.05 for 4% DSS). Histological analysis (2X magnification) indicates reduced loss of colon crypt architecture as seen on representative micrographs (**F-I**, 20X magnification, **G** and **H** provide 2 examples of damaged crypt architecture and inflammatory cell invasion). Mean damage as measured by length of colon damage in 4 histology sections and normalized to the length of the colon section examined per mouse (2X magnification) demonstrates significantly reduced damage for PEGSerp-1 treated mice given 5% DSS (**J**, *p* < 0.011) or 4% DSS (**K**, *p* < 0.0037). Mean values were calculated for measured colon damage normalized to length examined for three colon sections per mouse. Red arrows – damaged residual crypts, Black arrows – inflammatory cell invasion, block arrow indicates preserved crypt structure.

Markers of disease severity (body weight, colon length and crypt morphology) were also examined for the acute prophylaxis groups. Body weight (Figure 2D), colon length measured on colon samples isolated after euthanasia (Figure 2E) and histological evidence for extent of colon damage as evidenced by loss of crypt architecture (Figure 2F-K) were assessed. Body weight, represented as a percentage of weight on day 0, is summarized in Figure 2D. There was significant weight loss for both 4 and 5% DSS colitis when compared to normal controls with no DSS induced colitis (p < 0.01 days 6-14). Due to the high mortality in the vehicle treated 5% DSS group, comparative body weight could only be assessed up to day 6 during treatments. However, in addition to the clear survival benefit, when comparing the weights recorded on day 9 for PEGSerp-1 treated mice after euthanasia, there was also a significant improvement in weight gain with PEGSerp-1 treatments by day 14 (mean weight PEGSerp-1 treated day 9 - 76g, day 14 - 84g; *p*< 0.0394). In the 4% DSS acute prophylaxis model, weight was significantly improved with PEGSerp-1 prophylaxis on days 7 to 10 (Figure 2D, *p* < 0.05). PEGSerp-1 treatment in the mice surviving the 4 and 5% DSS colitis treatments demonstrated ongoing improved weight gain on days 12 to 14, indicating that general colon health continued to improve after discontinuing DSS and with PEGSerp-1 treatment, demonstrating a potential impact on recovery (Figure 2D).

Colon shortening is considered a marker for colitis severity in the DSS models and is used as a marker of disease progression and tissue damage. Colon length measured immediately after euthanasia on excised colon sections was significantly increased in PEGSerp-1 treated mice (Figure 2E*, p* < 0.0469) compared to saline vehicle treatment in the acute prophylaxis 4% DSS group. Within the acute prophylaxis 5% DSS group, PEGSerp-1 treatment produced a trend toward reduced colon shortening (Figure 2E, *p* = 0.1921). This was not statistically significant (Figure 2E, *p* = 0.1921), which may be attributed to the early mortality in the 5% saline control group. Combining colon length measurements from 4% and 5% DSS induced colitis, however, did demonstrate a significant improvement for the acute severe DSS colitis models, approximating more normal colon lengths in mice without DSS treatments (Figure 2E). Enlarged spleens were also observed in this acute prophylaxis model in Saline controls with reduced spleen size noted on visual inspection in the PEGSerp-1 treated mice, but changes in spleen size were not quantified in these first experiments.

Colon sections were also examined by blinded histopathological analysis for disease severity, specifically loss of architecture (Figure 2J, K). Histology sections from colon samples were analyzed for classical histological changes associated with colitis damage including loss of architecture and decreased crypt density, crypt structural distortion and irregular mucosal surface. In both the 4% and the 5% DSS groups, acute prophylactic treatment with PEGSerp-1 significantly reduced the length of damage (loss of crypt architecture) relative to total length of colon section assessed, as measured by blinded analysis on histologic sections from the 5% (Figure 2F-J; *p* < 0.0011) and 4% DSS models (Fig 2K; *p* < 0.0037).

### Acute and chronic delayed PEGSerp-1 treatment in 2%DSS colitis models

The acute colitis induced by 4 and 5% DSS indicated potential benefit for early PEGSerp-1 treatments in severe colitis, with improved mortality and reduced colon damage when PEGSerp-1 treatment was given prophylactically at the time of inducing colitis (Figure 1A). In clinical settings it is more common to see chronic colitis and start treatments at delayed times after initial diagnosis or during episodes of increased disease activity, *e.g.* acute exacerbations. In order to examine the potential for benefit with delayed PEGSerp-1 treatment, two studies were next performed using a less aggressive 2% DSS induced colitis model (Figure 1B, C), designed to simulate less aggressive, chronic colitis with delayed treatments, as seen in clinic. In the first study, 2%DSS was given to induce gut inflammation and PEGSerp-1 treatment was not started until 7 days after first 2% DSS challenge, termed acute delayed treatment (Figure 1B). In the chronic delayed treatment study, the third model, a chronic relapsing colitis was simulated where 2% DSS was given for 5 days followed by 5 days of normal water and then repeat introduction of 2% DSS as a second DSS bolus starting on day 10 to simulate a recurrent or acute exacerbation of colitis (Figure 1C). PEGSerp-1 treatment was started on day 10 and continued for 3 days (total of 4 days) to simulate a treatment for acute recurrent colitis exacerbation, termed chronic delayed treatment for chronic colitis.

No significant improvement in survival was seen with acute delayed PEGSerp-1 treatment starting 7 days post 2% DSS challenge (Figure 3A), nor with chronic delayed PEGSerp-1 treatment given with the re-introduction of 2%DSS at 10 days (Figure 3B). In the acute delayed treatment study, four of the saline vehicle treated mice died at 8-10 days, 1 to 3 days after initiating saline treatment, whereas three of the PEGSerp-1 mice died at 9 days, 2 days after initiating PEGSerp-1 treatments. This trend did not reach significance (Figure 3A, *p* = 0.5488). In the chronic delayed treatment colitis, where PEGSerp-1 was started after re-introducing 2% DSS to the drinking water on day 10, there was again no significant improvement in mortality (Figure 3B, *p* = 0.4747). Body weights were also not significantly different for saline vehicle and PEGSerp-1 treatments in the acute colitis with delayed treatment group (Figure 3C, *p* = ns at all times), nor in the chronic delayed treatments (Figure 3D at all times). However, weight did gradually improve in all treatment groups, whether acute delayed or chronic delayed treatments indicating potential for recovery from the damage to the colon and no interference with improved weight gain with treatment.

**Figure 3.**
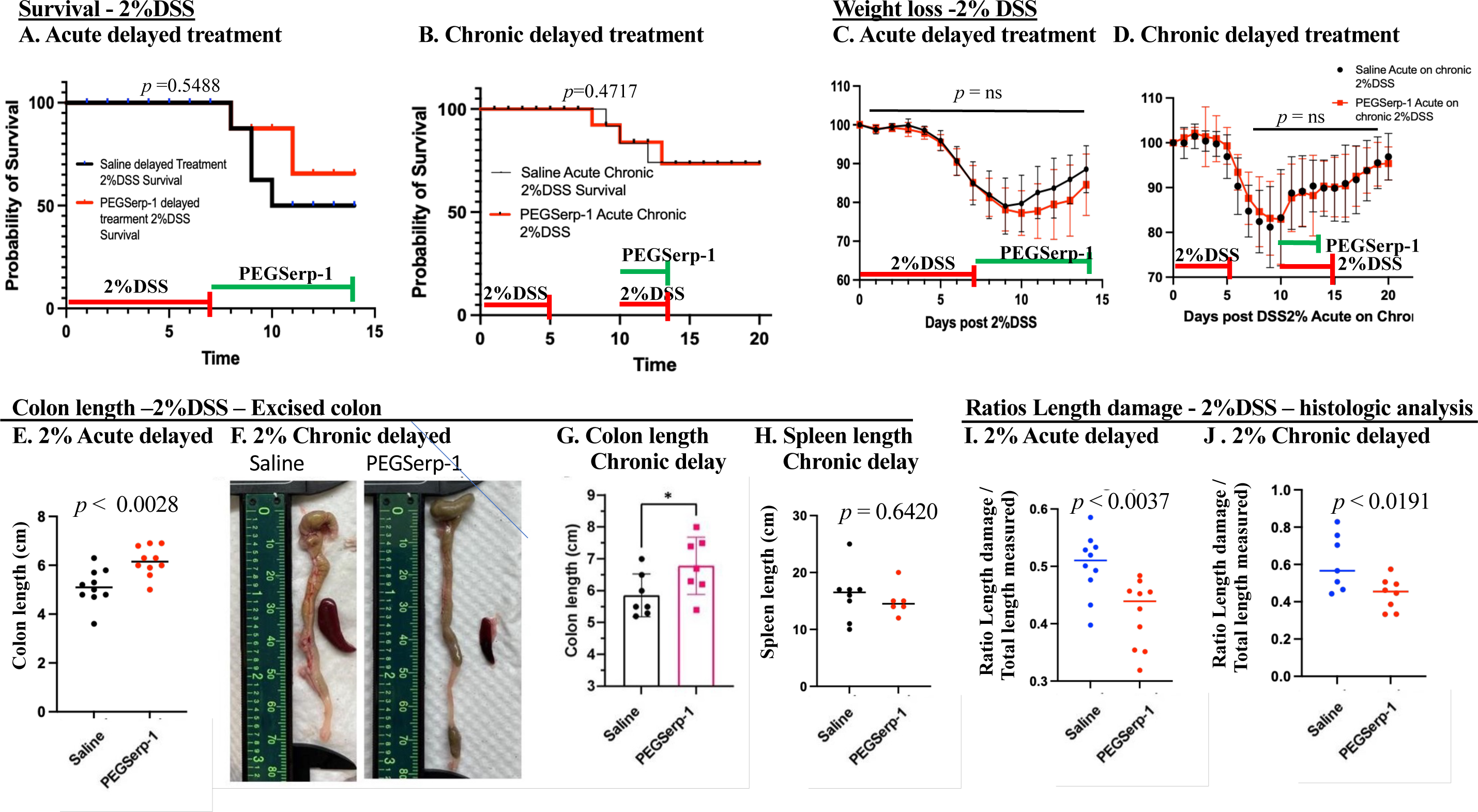
2%DSS induced colitis with either acute delayed PEGSerp-1 treatment starting 7 days after introducing DSS to the drinking water (**A**, *p* = 0.5488) or when given on introducing a second round of 2%DSS in the drinking water 10 days after the initial 5 days of 2% DSS (**B**, *p* = 0.4717) did not improve survival. Timing for DSS introduction is indicated in red and for PEGSerp-1 treatment is indicated in green at the base of the graphs. Wight loss was also not significantly altered by PEGSerp-1 treatments although in the acute delayed (C, p = ns) and the chronic delayed treatments (D, *p* = ns). There is a gradual improvement in weight gain after discontinuing DSS demonstrating the capacity for recovery. Excised colons removed after euthanasia demonstrate increased colon length in the Acute delayed treatment model (**E**, *p* < 0.0028). Measurement of colon length is illustrated for the 2%DSS chronic delayed treatment model (**F**) and demonstrates increased colon length with PEGSerp-1 treatments (**G**, *p* < 0.05). Spleen size appeared larger in the saline treated group but did not achieve significance on measured length (**H**, *p* = 0.6420). Measured damage length normalized to colon section length examined on histological analysis detected significantly reduced damage with the acute delayed (**I**, *p* < 0.0037) and the chronic delayed (**J,** *p* < 0.0191) treatments with PEGSerp-1. Mean values were calculated for measured colon damage normalized to length examined for three colon sections per mouse.

Colon length, measured immediately after euthanasia and excision, was significantly increased with PEGSerp-1 treatment in the acute delayed (Figure 3E, *p* < 0.0028) and in the chronic delayed treatment groups (Figure 3F, G; *p* < 0.05). It has been reported that increased spleen size generally correlates with the severity of inflammation (36,37). For mice that reached the end of the study, 6/7 saline treated mice had visibly enlarged spleens compared to 2/6 mice in the PEGSerp-1 acute delayed treatment group (*p* < 0.05, Chi square analysis). In the chronic delayed treatment group, a reduction in spleen size was also noted by visual inspection but measured spleen length was not significantly reduced (Figure 3H, *p* = 0.6420). Similar to the acute prophylaxis treatments in the 4% DSS and 5% DSS colitis models, blinded histological analysis further confirmed reduced architectural damage for acute delayed PEGSerp-1 treatment starting on Day 7 after 7 days of 2% DSS induced colitis (Figure 3I, *p* < 0.0037) and for the chronic delayed treatment, where 2% DSS was re-introduced at 10 days together with initiating delayed PEGSerp-1 treatments (Figure 3J, *p* < 0.0191).

### PEGSerp-1 treatment reduced inflammatory cell invasion and loss of crypt architecture

The depth of inflammatory cell infiltrates together with loss of crypt length was further assessed in the DSS induced colitis models, the acute prophylaxis in 4% and 5% DSS model as well as in the acute and chronic delayed 2% DSS models. Depth of transmucosal inflammatory cell infiltrates as well as preserved crypt architecture (crypt length) were measured on colon histology sections and normalized to colon wall thickness.

For mice in the acute prophylaxis 4% and 5% DSS groups, histological assessment demonstrated a reduction in the depth of immune cell infiltrates in those treated with PEGSerp-1 (Figure 4). The depth of inflammatory cell infiltrates, normalized to the mucosal wall thickness, was significantly reduced by prophylactic PEGSerp-1 treatment in the acute severe 5% DSS (Figure 4A, *p* < 0.0013) and the 4% DSS groups (Figure 4C, *p* < 0.0086). Crypt depth, normalized to colon wall thickness, was also used to assess colon damage and preservation of normal architecture. There was a trend toward increased crypt depth in the acute prophylaxis 5% DSS model (Figure 4B, *p* = 0.1712) and a significant increase in crypt depth normalized to mucosal wall thickness in the 4% DSS acute prophylaxis model (Fig 4D, p < 0.0396). Representative histology cross sections for normal colon (no DSS) and Saline treated and PEGSerp-1 treated colon are provided in Figure 4F. Combined analysis of 4% and 5% analyses for inflammatory cell invasion as well as preserved crypt depth was similarly significant (Figure 4E, G, *p* < 0.0001 and *p* < 0.0253, respectively). In the acute delayed 2%DSS model, the depth of immune cell infiltrates (Figure 4H, *p* = 0.2308) and the crypt depth (Figure 4I, *p* = 0.4291), normalized to the colon wall, were not significantly modified by PEGSerp-1 treatments. In contrast, in the chronic delayed 2% DSS model PEGSerp-1 treatment significantly reduced the depth of inflammatory cell infiltrates into the colon wall (Figure 4J, *p* < 0.0222) and significantly increased crypt depth (Figure 2K, *p* < 0.0015). Blinded histopathological analysis of the 5% and 4% DSS acute prophylaxis histology sections (SG) confirmed a significant reduction in inflammation and damage (Figure 4L, p < 0.0108).

**Figure 4.**
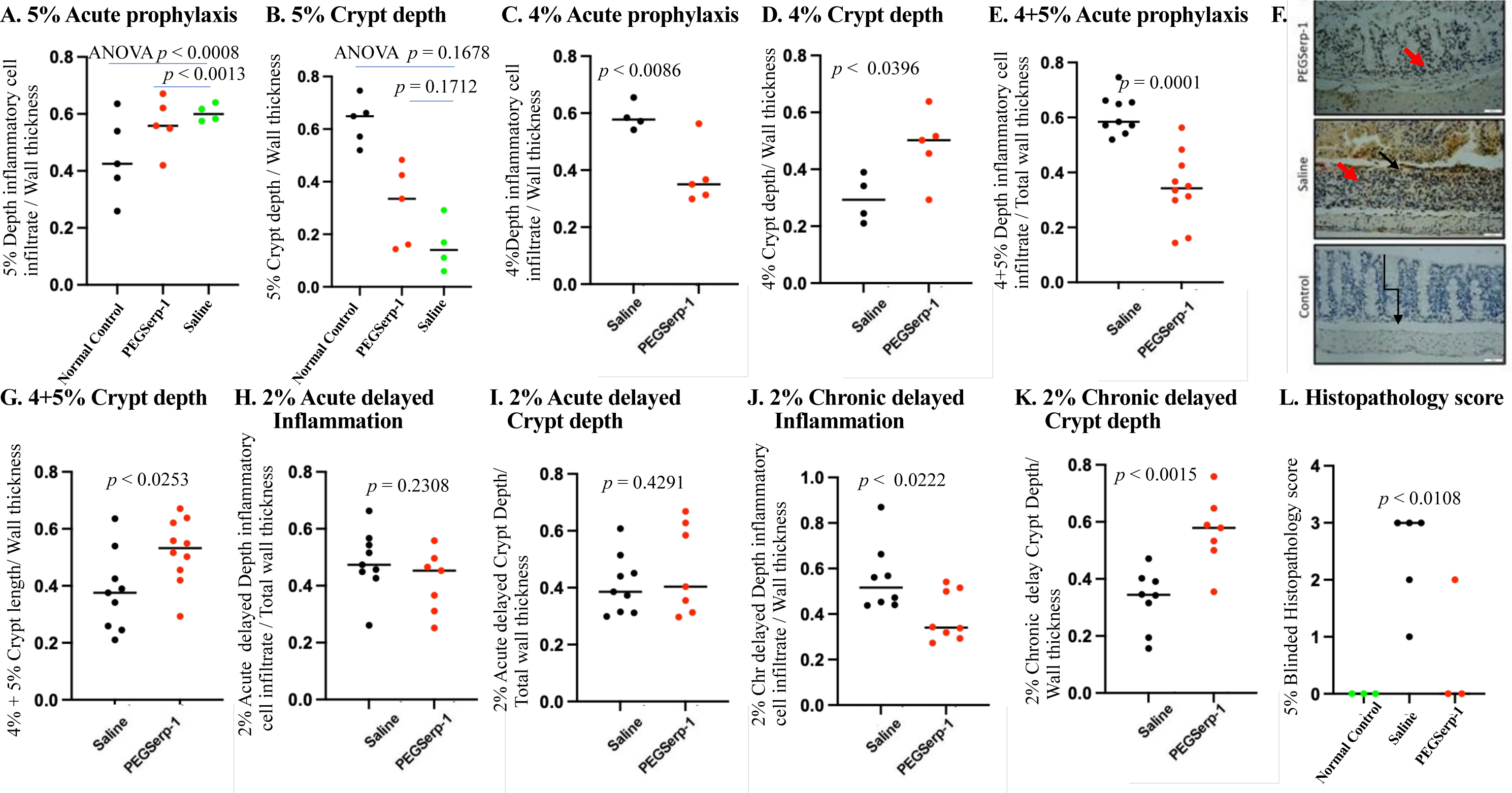
PEGSerp-1 treatment reduced the depth of inflammatory cell infiltrates and improved crypt architecture as measured by crypt depth, measurements normalized to total colon diameter. Inflammatory cell infiltrate depths divided by total colon thickness is reduced (**A**, *p* < 0.0013) and crypt depth divided by total colon thickness is increased (**B**, non-significant trend *p* = 0.1712) with acute PEGSerp-1 treatment with 5% DSS induced colitis. With 4% DSS colitis PEGSerp-1 reduced the inflammatory infiltrate (**C**, *p* < 0.0086) and increased crypt depth (**D**, p < 0.0396), each normalized to wall thickness. Combined analysis of data for acute PEGSerp-1 prophylaxis for 5% and 4% DSS induced colitis again indicated significant reduction in inflammatory cell invasion (E, *p* < 0.0001) and improved crypt architecture as measured by crypt depth (G, *p* <0.0253), each normalized to colon wall thickness. Histology sections from the 5% DSS acute prophylaxis treatment group illustrate extensive inflammatory cell invasion with loss of crypt depth in saline treated mice with improved architecture and reduced inflammation in the PEGSerp-1 treated mice (F). PEGSerp-1 treated sections have preserved architecture closer to normal mouse colon with no DSS induced colitis (F). Acute delayed PEGSerp-1 treatment in the 2% DSS model did not have significantly reduced cell infiltrates (**H**, p= 0.2308) nor increased crypt depth (**I**, p = 0.4291) normalized to colon wall thickness. Chronic delayed PEGSerp-1 treatment in the 2%Dss model, initiated at 10 days when 2% DSS was re-introduced to the drinking water simulating acute exacerbation did significantly reduce inflammation **(J**, *p* < 0.0222) and improve crypt architecture (**K**, *p* < 0.0.0015). A blinded histopathological score demonstrated significantly reduced scores for colitis in the 5% and 4% DSS histology section with acute prophylaxis PEG Serp-1 treatments (**L**, *p* < 0.0108).

### PEGSerp-1 treatment reduces inflammatory macrophage invasion in DSS-induced colitis

A loss of balance in macrophage and T cell responses that disturbs epithelial integrity is closely associated with ongoing colon damage in IBD. Inflammatory cell invasion was further assessed in each DSS colitis model by immunohistochemical analysis of immune cell infiltrates, specifically iNOS+ M1 macrophage, Arg 1+ M2 macrophage, Ly6G+ neutrophils, and CD3+ and CD 4+ T lymphocytes. PEGSerp-1 has been previously demonstrated to reduce proinflammatory M1 macrophage invasion in animal models of inflammatory disease and in some cases to increase Arg-1+ M2 cell infiltrates (38,39). Modified T cell responses have also been detected in select models with Serp-1 treatment (39,40).

IHC analysis of iNOS+ M1 pro inflammatory macrophage cell infiltrates detected a borderline reduced M1 macrophage count in the acute prophylaxis 5% DSS model (Figure 5A and B, *p* < 0.0587). iNOS cell counts were also reduced, but again not significantly in the acute prophylaxis 4% DSS (Figure 5C, *p* = 0.6765) colitis. Combined analysis of 4% and 5% iNOS+ M1 counts indicates significantly reduced iNOS M1 macrophage counts (Figure 5D, *p* < 0.0278). M1 macrophage count were also significantly reduced with PEGSerp-1 treatment, both in the acute delayed 2%DSS (Figure 5E, *p* < 0.0454) and the chronic delayed 2% DSS treatments (Figure 5F, *p* < 0.0005) models. M2 Arg1+ anti-inflammatory macrophage counts were not significantly modified by PEGSerp1-treatments in any of the models examined (Figure 5 G-J, *p* = ns).

**Figure 5.**
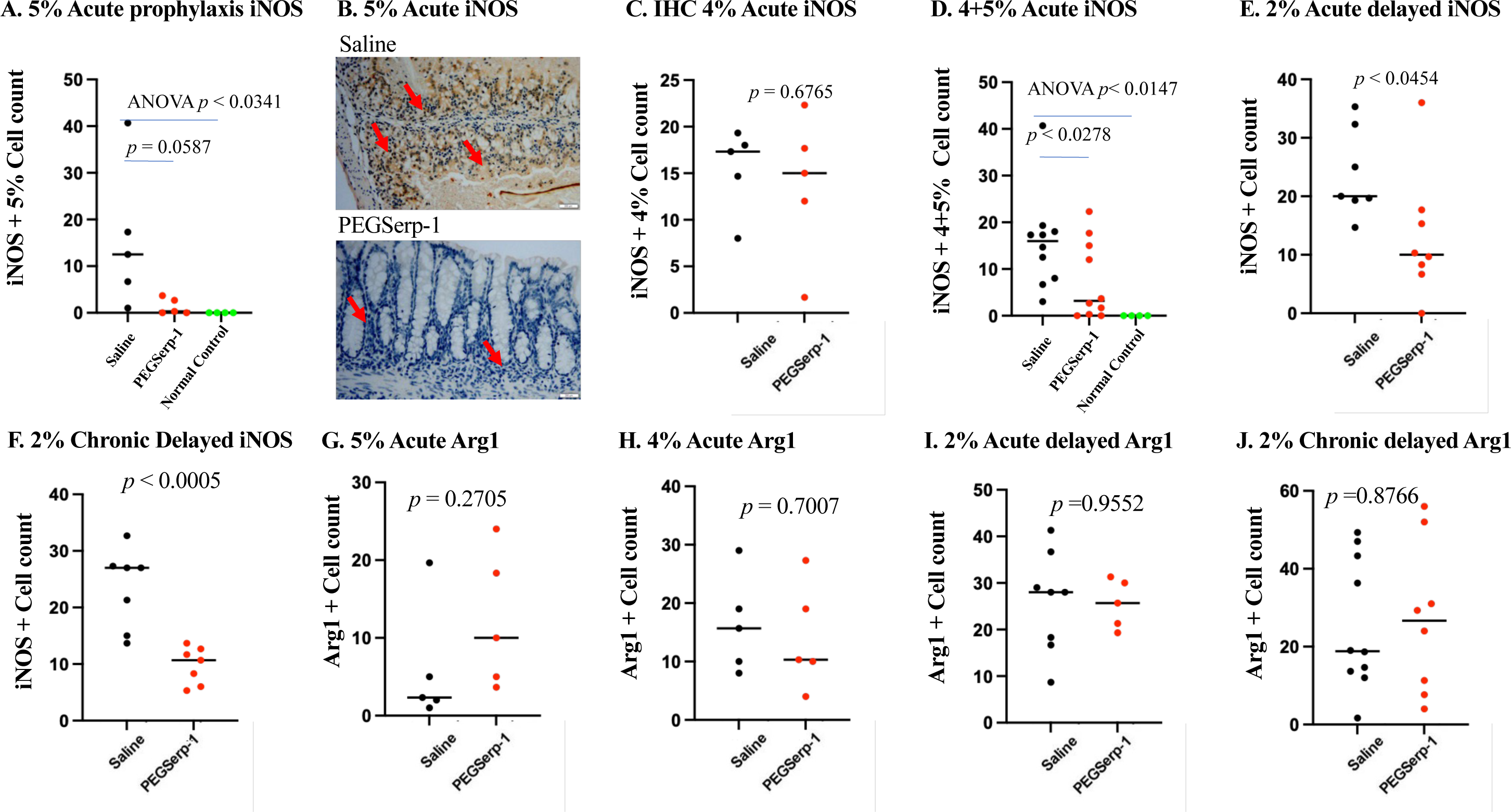
PEGSerp-1 treatment given as acute prophylaxis or as a chronic delated treatment significantly reduced iNOS positive M1 macrophage infiltrates in DSS colitis. Significantly reduced mean iNOS counts in 5% DSS with Acute prophylaxis PEGSerp-1 treatment (**A**, ANOVA *p* < 0.0341, borderline *p* < 0.0587 Fishers PLSD for PEGSerp-1 vs Saline). iNOS stained 5% DSS histology sections illustrate loss of crypt architecture and extensive iNOS positive cell infiltrates with Saline treatments (**B**). There is reduced iNOS staining and improved crypt architecture with PEGSerp-1 treatment (**B**). Acute prophylaxis with PEGSerp-1 with 4% DSS did not significantly reduce iNOS staining (**C**, *p* = 0.6765), however combined analysis of 4 and 5% PEGSerp-1 acute prophylaxis detected significantly reduced iNOS positive cell infiltrates (D, *p* < 0.0278). iNOS positive cell counts were significantly reduced with PEGSerp-1 treatment in both the 2%DSS Acute delayed (E, *p* < 0.0454) and the Chronic delayed treatment models (**F**, p < 0.0005). Arg1+ M2 macrophage staining was not significantly modified by PEGSerp-1 in the 5% (**G**, *p* = 0.2705), 4% (**H**, *p* = 0.7007), nor the 2% acute (**I**, p = 0.9552) and chronic delayed DSS colitis models (**J**, p = 0.8766). IHC analysis performed at 20X, mean values calculated for 3 sections with visible positive staining. Micrographs – 20X. Red arrows indicate iNOS positive cells – IHC analysis.

In addition to macrophage invasion, Ly6G+ neutrophil and also CD3+ and CD4+ T lymphocyte activation have also been closely associated with active colitis. No significant changes in Ly6G+ neutrophil counts were detected in the acute prophylaxis 5% (Figure 6A, *p* = 0.6918) and 4% DSS models (Figure 6B, *p* = 0.79981) with PEGSerp-1 treatments, nor in the chronic delayed treatment model (Figure 6D, *p* = 0.2640). Ly6G counts were reduced in the 2% acute delayed treatment 2% DSS model (Figure 6C, *p* < 0.05). CD3 positive T cell counts were not modified with acute prophylactic PEGSerp-1 treatment in the 5 and 4% DSS models (Figure 6E and F, *p* = 0.5757 and *p* = 0.3894, respectively), but had a nonsignificant trend toward a reduced count in the 2% DSS acute delayed treatment model (Figure 6G, *p* = 0.0869) with PEGSerp-1 treatment. PEGSerp-1 treatment also did not alter detectable CD4+ T lymphocyte counts in the acute prophylaxis 5% and 4% DSS models (Figure 6H, I, *p* = 0.2556 and *p* = 0.4481, respectively), nor in the acute delayed 2% DSS treatment model (Figure 6J, *p* = 0.0628). In contrast, in the chronic delayed 2% DSS model there is a significant reduction in CD4 + T cell counts with PEGSerp-1 treatment (Figure K, *p* < 0.0016).

**Figure 6.**
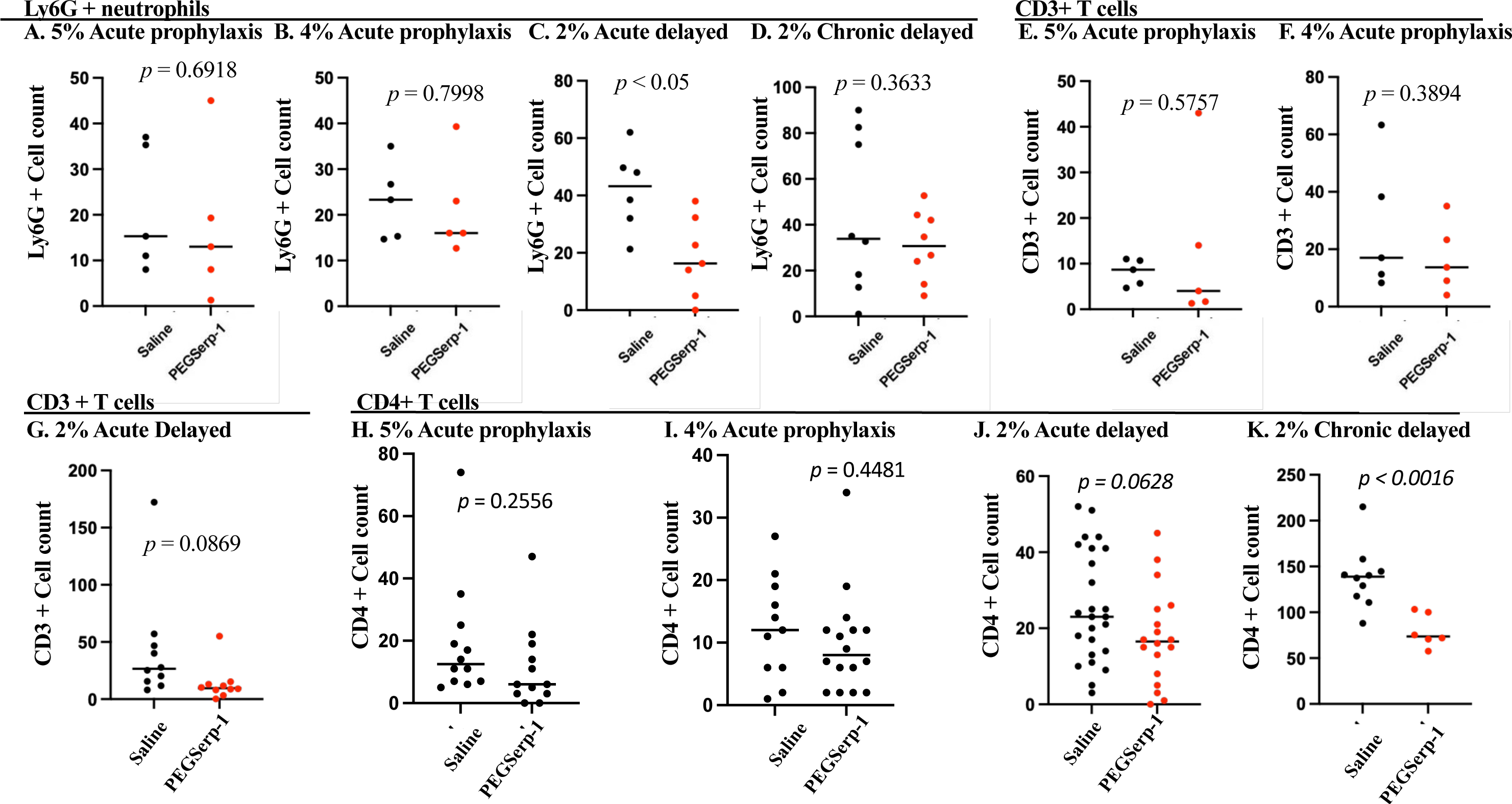
PEGSerp-1 acute prophylaxis treatments in the 5% (**A,** *p* = 0.6918) and 4% (**B,** *p* = 0.7998) DSS colitis models, and the 2% chronic delayed (**D,** *p* =) colitis models had no significant effect on Ly6G + neutrophil cell counts. In the acute delayes 2% DSS model, PEGSerp-1 tretament did significantly reduced Ly6G+ cell counts (**C,** *p* < 0.05) Nonspecific CD3+ T cell counts were similarly unaffected by PEGSerp-1 treatments in the 5% (**E**, *p* = 0.5757) and 4% (**F**, *p* = 0.3894) acute prophylaxis and the 2% acute delayed colitis models (**G**, *p* = 0.0869). CD4 + T cell counts were not modified by PEGSerp-1 treatment in the 5% (**H**, *p* = 0.2556) and 4% (**I**, *p* = 0.4481) acute prophylaxis models and in the 2% (**J,** *p* = 0.0628) acute delayed model. However, CD4+ cell counts were significantly reduced with PEGSerp-1 treatment in the 2% chronic delayed colitis treatment model (**K**, *p* < 0.0016).

Overall, these studies indicate that PEGSerp-1 treatment reduces inflammation and crypt damage when given early as an acute prophylaxis in severe 4% and 5% DSS induced colitis, as well as when given as an acute delayed treatment or a chronic delayed treatment with 2% DSS induced colitis. This reduced inflammation was greater for the 4% and 5% DSS acute prophylaxis and the 2% DSS chronic delayed models when compared to the 2% DSS acute delayed treatment model. In the latter model, PEGSerp-1 treatment given 7 days after initiating colitis with 2% DSS, acute delayed treatment, had less impact on overall inflammation. M1 iNOS+ pro-inflammatory macrophage infiltrates were significantly reduced in each model system whereas altered Ly6G neutrophil counts were not modified on IHC analysis. CD3+ T cells were not changed with PEGSerp-1 treatment. CD4+ cell infiltrates were reduced, but only reaching significance in the chronic delayed 2% DSS model consistent with a late effect on T cell response and is also consistent with the known timing predicted for early macrophage and later T cell responses.

### PEGSerp-1 treatment significantly alters detected coagulation pathway factors on IHC analysis

The coagulation pathways, both thrombotic and thrombolytic, are closely associated with inflammatory responses. Serp-1 binds fXa and thrombin in the clotting cascade as well as uPA and its receptor (uPAR), in the fibrinolytic pathway. In the clotting cascade, fibrinogen (Figure 7 A-G) and fXa (Figure 7H-N) were assessed by IHC in all DSS colitis models by IHC analysis. MAC were also assessed.

**Figure 7.**
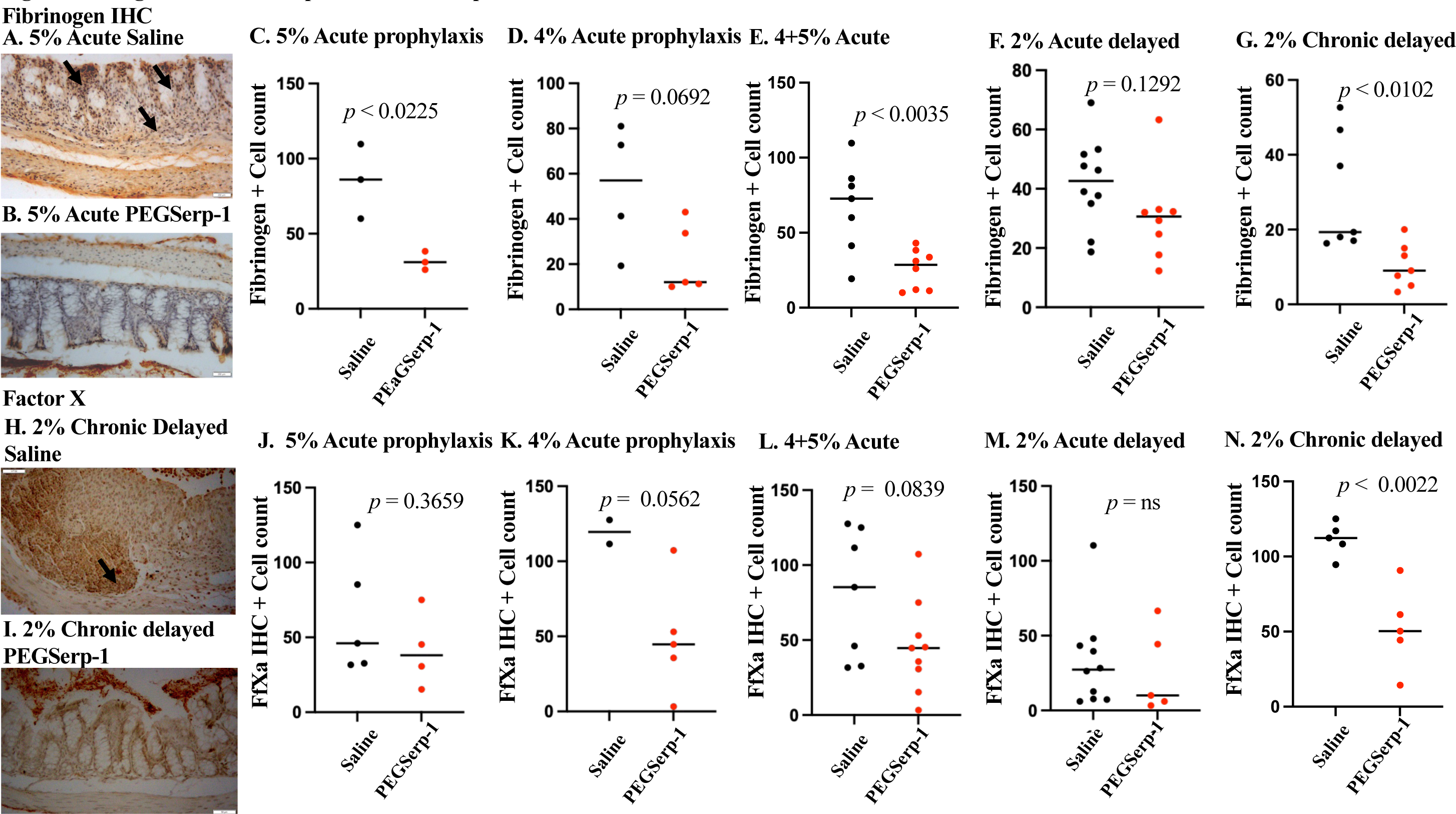
IHC stained sections from 5% DSS acute prophylaxis colitis model (**A** - Saline, **B** - PEGSerp-1 acute prophylaxis) illustrate reduced fibrinogen positive staining and improved colon, crypt architecture. Fibrinogen positive counts were significantly reduced with acute prophylaxis in the 5% DSS (**C**, *p* < 0.0225), but not the 4% DSS albeit with a trend toward a reduction (**D**, *p* = 0.0692). Combined 4% and 5% DSS counts were significantly reduced with PEGSerp-1 treatments (**E**, *p* < 0.0035). In the 2% DSS acute delayed (colitis model there was no significant reduction in fibrinogen + cells (**G** *p* = 0.1292). Fibrinogen positive staining on IHC colon sections illustrates significantly reduced in the chronic delayed 2% colitis model (H - Saline, I-PEGSerp-1). FXa IHC staining detected no significant reduction in positive staining with PEGSerp-1 treatment in the 5% DSS (K, *p* = 0.3659), but borderline significant reduction in the 4% DSS (K, *p* < 0.0562) and in combined 4% and 5% fXa staining (L, p = 0.0839). In the 2% acute delayed colitis treatment model fXa was not reduced with acute delayed PEGSerp-1 (**K**, *p* = ns) treatment, but was significantly reduced with 2% chronic delayed PEGSerp-1 treatment (L, *p* < 0.0022). Black arrows – fibrinogen and fXa positive stained areas. Mag 20X.

Detectable fibrinogen staining was significantly reduced in the 5% DSS colitis models with acute PEGSerp-1 prophylaxis (Figure 7A-C, 5% DSS p < 0.0225) while PEGSerp-1 prophylaxis for the 4% DSS challenge only produced a trend toward a reduction in fibrinogen staining (Figure 7D, *p* = 0.0625). Combined 4% and 5% acute DSS groups did show a significant reduction in fibrinogen staining on IHC with PEGSerp-1 treatment (Figure 7E, *p* < 0.0035). Fibrinogen staining was not significantly reduced in the 2% acute delayed DSS model (Figure 7F, *p* = 0.1292), but was significantly reduced with PEGSerp-1 treatment in the 2% chronic delayed DSS colitis model (Figure 7G, *p* < 0.0102).

Detected fXa staining was also reduced on IHC analysis in the 5% and 4% DSS acute prophylaxis models, but not significantly (Figure 7J,K, *p* = 0.3659 and *p* = 0.0562, respectively), albeit a trend for 4% DSS is seen and with combined 4% and 5% DSS group analysis comparing PEGSerp-1 to Saline vehicle treatment (Figure 7L, *p* < 0.0839). FXa staining was not reduced in the 2% DSS acute delayed treatment group (Figure 7M, *p =* ns), but was significantly reduced in the 2% DSS chronic delayed treatment group (Figure 7H, I, N; *p* < 0.0022) with PEGSerp-1 treatment.

### PEGSerp-1 modifies detectable inflammatory uPAR and complement C5b/9 MAC levels

Serp-1 binds uPA, uPAR and proteases in the complement pathway. uPAR is considered a predominantly inflammatory mediator, forming a complex with uPA that activates MMPs allowing immune cell invasion and is reported to be upregulated in IBD (10) while C5b/9 is the complement MAC that effectuates innate and acquired immune cell responses. Both components were assessed by IHC analysis in the three DSS colitis models.

uPAR+ cell counts were significantly increased when compared to normal colon (Figure 8C, ANOVA *p* < 0.0365), but were not significantly reduced by PEGSerp-1 treatment in the acute prophylaxis 5% DSS model (Figure 8C, 5% DSS, *p* = 0.1688). In contrast, uPAR+ counts were significantly reduced in the acute 4% DSS (Figure 8D, *p* < 0.0237) model with PEGSerp-1 treatment. Combined analysis of uPAR detection for 4% and 5% DSS acute prophylaxis indicated a trend toward reduced uPAR+ (Figure 8E, *p* < 0.0610). No significant reduction in uPAR observed in the 2% acute delayed colitis model (Figure 8F, *p* = 0.3433), but this was seen for the chronic delayed 2% DSS acute group (Figure 8A, B, G, *p* < 0.0158).

**Figure 8.**
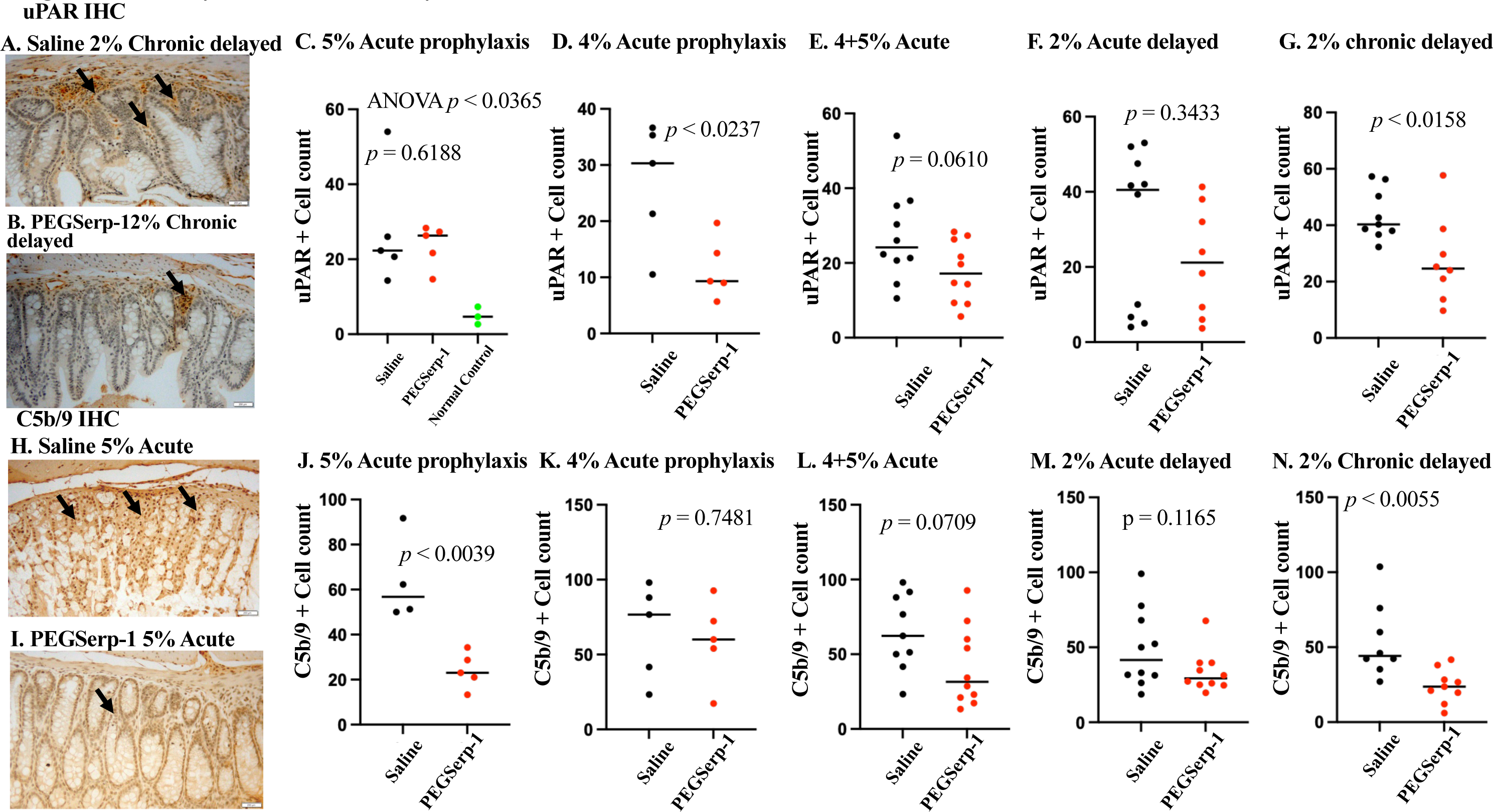
uPAR and C5b/9 positive staining was significantly reduced with acute prophylaxis 4 and 5% DSS models with PEGSerp-1 treatment. Micrographs illustrate IHC analysis demonstrating reduced uPAR for 2% DSS chronic delayed treatment (**A**-Saline, **B** - PEGSerp-1). uPAR was not reduced for the 5% acute prophylaxis treatment with PEGSerp-1 (**C**, *p* = 0.6188) but was significantly reduced for the 4% DSS acute prophylaxis (**D**, *p* < 0.0237). Combined 4 and 5% DSS acute prophylaxis indicates a trend toward reduced uPAR (**E**, p = 0.0610). In the 2% acute delayed treatment there was no significant reduction in uPAR staining (**F,** *p* = 0.3433), but for chronic delayed treatment for 2% DSS colitis PEGSerp-1 did significantly reduce uPAR detection (**G**, *p* < 0.0158). Complement C5b/9 + staining is reduced with improved crypt structure (H – Saline, I - PEGSerp-1) in 5% DSS acute prophylaxis with PEGSerp-1. There is a significant reduction with PEGSerp-1 treatments in 5% DSS prophylaxis (**J**, *p* < 0.0039), but not 4% DSS (**K**, *p* = 0.7481). Combined 4% and 5% acute prophylaxis indicates a trend toward a reduction (**L**, *p* = 0.0709). 2% Acute delayed PEGSerp-1 did not reduce detected C5b/9 staining (**M**, *p* = 0.1165), but PEGSerp-1 did significantly reduce C5b/9 detection in the 2% chronic delayed colitis model (**N**, *p* < 0.0055). Black arrows indicate cells positively stained for uPAR or C5b/9 on IHC analysis. Mag 20X.

PEGSerp-1 also showed a trend towards reduced C5b/9 detection, with a significant reduction seen in the 5% acute prophylaxis colitis model (Figure 8H-J, *p* < 0.0039). PEGSerp-1 treatment did not reduce detected C5b/9 in the 4% DSS acute model (Figure 8K, *p* = 0.7481), nor the acute delayed 2% DSS model (Figure 8M, *p* = 0.1165). Combined IHC data analyses for the 4% and 5% DSS acute prophylaxis DSS colitis models indicated a trend toward a reduction with PEGSerp-1 treatments (Figure 8L, *p* = 0.0709). In the chronic delayed 2% DSS colitis model, PEGSerp-1 treatments again significantly reduced C5b/9 detection (Figure 8N, *p* <. 0.0055).

### PEGSerp-1 treatment reduces inflammatory responses in submucosal colon vasculature

Based upon the known close association between IBD colon inflammation and inflammatory systemic cardiovascular disease, venous thrombosis, and local colon micro-clots with gut ischemia, vascular inflammation and coagulation were assessed in the submucosal vessels in the acute prophylaxis 5% DSS colitis model and in the 2% DSS chronic delayed PEGSerp-1 treatment models. In prior work, treatment with Serp-1 reduced inflammatory plaque in animal models of arterial injury including balloon angioplasty, hyperlipidemic plaque, transplant vasculopathy and virus induced vasculitis (gamma Herpes and SARS MA30 viral infection models) (17,30–32). These observed reductions in vascular inflammatory cell infiltrates were associated with reduced iNOS positive M1 macrophage and in some cases also reduced CD4+T cells. Serp-1 treatment also reduced markers of heart damage after coronary stent implant (35). These inflammatory changes in prior models were linked to altered uPAR and C5b/9. To assess the mechanisms for reduced colon inflammation and damage with PEGSerp-1 treatments, we examined smaller vessels in the colon submucosal and adventitial layers associated with colitis. The analysis of vascular inflammation and dysfunctional coagulation was focused on the chronic delayed 2% DSS model which more closely simulates acute recurrent episodes, acute exacerbation of colitis observed in the clinic.

Significantly reduced iNOS+ M1 macrophage staining was detected in the submucosal vasculature in the colon with prophylactic PEGSerp-1 treatments in the acute 5% DSS colitis model (Figure 9, Panels 1A-C, *p* < 0.0481), but was no longer significantly decreased in the 2% DSS chronic delayed colitis PEGSerp-1 treatment model (Figure 9, Panel 1D, *p* = 0.2989). Arg1+ M2 macrophage and CD4+ T cell counts were also not modified in the 2% DSS chronic delayed PEGSerp-1 treatment model (Figure 9, Panels 1E-F, *p* = 0.5915 and *p* = 0.1110, respectively), although CD4+ T cell counts showed a negative trend. In contrast, analysis of the clotting pathway factors demonstrated a significantly reduced detectable staining for fXa (Figure 9, Panel 1G-I, p < 0.0331) and for fibrinogen (Figure 9, Panel 1J, p < 0.0502) with PEGSerp-1 treatment in this model. Although uPAR detection was significantly decreased in the colon mucosal layers, uPAR detection was not reduced in the submucosal vessels in the acute 5% DSS (Figure 9, Panel 1K, *p* = 0.1476) with PEGSerp-1 treatment, nor in the chronic delayed 2% DSS colitis models (Figure 9, Panel 1L, *p* = 0.1675). IHC detection for complement (C5b/9) positive staining was reduced with borderline significance in vessel walls in the 2% chronic delayed colitis model with PEGSerp-1 treatment (Figure 9, Panel 1M, *p* = 0.0571).

**Figure 9.**
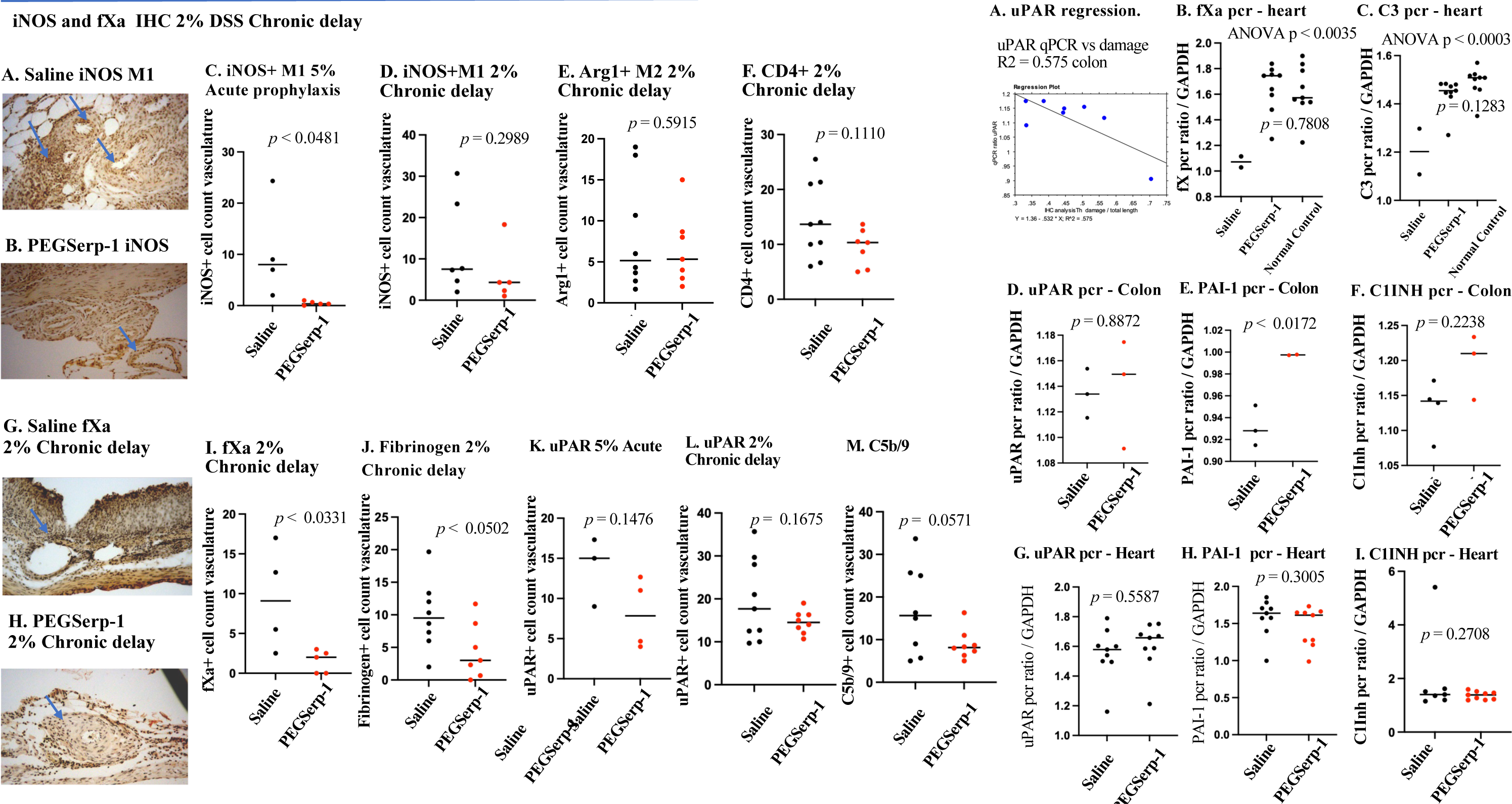
Panel 1 - Micrographs illustrate reduced iNOS+ M1 macrophage positive staining in the submucosal vessels (**A**-Saline, **B**-PEGSerp-1). iNOS positive counts are significantly reduced in the 5% DSS colitis model (**C**, *p* < 0.0481), but not in the 2% chronic delayed treatment colitis model (**D**, *p* = 0.5915). Arg1+ M2 macrophage positive cell counts (**E**, p = 0.5915) and CD4+ T cell counts (**F,** p = 0.1110) in the submucosal vessels were not altered in the 2% chronic delayed model. fXa+ cells (**G-I**, *p* < 0.0331) and fibrinogen+ cells (**J,** *p* < 0.0503) were significantly reduced in the 2% chronic delayed treatment model. uPAR positive staining was not reduced with PEGSerp-1 in the 5% acute prophylaxis (K, *p* = 0.1476) nor the 2% chronic delayed (L, *p* = 0.1675) models. C5b/9 positive cells in the 2% DSS Chronic delayed colitis model were borderline significantly reduced in submucosal vessels in the 2%chronic delayed colitis model with PEGSerp-1 treatment (**M**, *p* = 0.0571). Dense perivascular aggregates of positively stained cells indicated by blue arrows. Mag 20X. Panel 2 – RT-PCR analysis of gene expression changes. uPAR expression significantly correlated with colon damage measured in the 2%DSS chronic delayed colitis model. Gene expression for fXa (B, p < 0.0035 ANOVA) and C3 (C, p < 0.0003ANOVA) were significantly increased in the cardiac sample extracts from the 2% DSS Chronic delayed model when compared to normal controls without DSS treatments. In the colon samples uPAR expression was increased, but not significantly (D, *p* = 08872). PAI-1 was significantly increased (E, p < 0.0172), and C1Inh showed a trend toward increased expression (F, *p* = 0.2238). In the heart no significant changes were detected for uPAR (G, *p* = 0.5587), PAI-1 (H, *p* = 0.3005)nor C1INH (I, *p* = 0.2708).

In summary, analysis of vascular responses to PEGSerp-1 treatment indicate a significant reduction in iNOS+ M1 on IHC analysis with 5% acute prophylaxis, but not 2% chronic delayed treatment. There was no detected significant change in vascular Arg1+ M2 macrophage and CD4+ T cells in the submucosal vasculature. A persistent reduction in clotting factors fXa and fibrinogen as well as complement MAC (C5b/9), but not uPAR, was detectable on IHC analysis of the submucosal vessels in the chronic delayed 2% DSS model with PEGSerp-1 treatment. This suggests a potential for greater persisting effects of PEGSerp-1 treatment in the clotting and complement pathways, than in the uPAR related inflammation pathway.

### qPCR analysis of inflammatory markers in heart and colon samples in the chronic delayed 2% DSS model

Targeting of these pathways by PEGSerp-1 treatment is further supported by altered expression and some similar trends in gene expression for uPAR, C1Inh and PAI-1 on qPCR analysis in colon and cardiac extracts in the chronic delayed 2% DSS treatment model. These changes are similar to changes in gene expression detected during PEGSerp-1 treatment in a mouse model of SARS-CoV-2 mouse model. There is a significant correlation in uPAR gene expression with measured colon damage and inflammation (Figure 9, Panel 2A, Simple regression, R2 = 0.575). However, overall greater change was seen in the colon samples by qPCR analysis than in the heart samples. Analysis of fXa (Figure 9, Panel 2B, p < 0.0035) and C3 (Figure 9, Panel 2, C, p < 0.0003) gene expression in the 2% DSS Chronic delayed treatment model detected increased gene expression in cardiovascular samples when compared to normal controls indicating a systemic inflammatory response to the DSS induced colitis. In the colon the majority of changes in gene expression after PEGSerp-1 treatment were non significant trends (Supplemental Figure S1) except for a significant increase in PAI-1 in the colon (Figure 9, Panel 2, D-F*, p* < 0.0172 for PAI-1 expression). As colitis is associated with a systemic increase in vascular inflammation gene expression was further examined in heart samples in the 2% DSS chronic delayed colitis model. uPAR (Figure 9, Panel 2G, *p* = 0.5587) gene expression showed a trend toward increased expression in heart samples with PEGSerp-1 treatment, whereas PAI-1 (Figure 9, Panel 2H, *p* = 0.3005) and C1INH (Figure 9, Panel 2I, *p* = 0.2708) gene expression were not altered. The remainder of the gene expression changes in the colon and heart extracts were not significant (Supplemental Figures S1 and S2, *p* = ns) and overall changes observed in the colon extracts appear greater than in the heart etracts. Some trends such as the decreased fXa gene expression were consistent with the findings of reduced detectable fXa staining on IHC analysis of colon and submucosal vessels.

## Discussion

Vascular inflammation and micro-thrombotic occlusions contribute to progressive colitis and colon damage in IBD (1, 2, 7–16, 19–21, 26–29). Serpins represent up to 10% of circulating proteins in mammals, regulating thrombotic, thrombolytic and complement pathways (17,18). Serpins of mammalian origin have been studied as potential therapeutic targets – or agents in their own right - for their role in inflammatory processes. Treatment with C1-inhibitor (C1Inh, SERPING1) reduced complement activation and is associated with reduced inflammation in colitis (14–16), whereas plasminogen activator inhibitor-1 (PAI-1, SERPINE1) blocks thrombolytic protease tissue type plasminogen activator (tPA) and increases inflammation in colitis (26). Mice deficient in PAI-1 and/ or mice treated with small molecule inhibitor of PAI-1 have reduced inflammatory colitis. Alpha 1 antitrypsin (AAT, SERPINA1) inhibits neutrophil elastase and inflammation, but is also reported to inhibit thrombotic proteases and treatment with AAT reduces colitis in mouse models (27,28). Antithrombin III (ATIII, SERPINC1) inhibits the thrombotic proteases and reduces inflammation in colitis (41).

Serp-1 is a virus-derived serpin with demonstrated systemic anti-inflammatory activities in a wide array of animal models of vascular diseases associated with immune and coagulation pathway dysfunction (17,30–35,38).It is able to bind and inhibit thrombolytic proteases tPA, uPA, and plasmin, thrombotic proteases fXa and thrombin, immune mediating uPA/uPAR complexes as well as complement proteases (23–25,31–33), sharing many targets with mammalian serpins that have demonstrated benefit in reducing colitis. A PEGylated form of this protein,

PEGSerp-1, has improved pharmacokinetic properties (33,34). PEGSerp-1 is a modified virus-derived serpin, with systemic anti-inflammatory activities in a range of animal models of vascular diseases associated with immune and coagulation pathway dysfunction (17, 33,34). PEGSerp-1 binds and inhibits serine proteases in the thrombotic, thrombolytic and complement pathways; pathways with bidirectional activation in IBD that increase colon and systemic vascular damage. Thrombotic, thrombolytic and inflammatory proteases drive colon inflammation and damage as well as vascular disease. PEGSerp-1 binds the thrombolytic pathway proteases tPA and uPA, the thrombotic proteases fX and thrombin, and immune mediating uPA/uPAR complexes and selective complement proteases (23–25,32–34). Systemic PEGSerp-1 treatment was evaluated here in DSS-induced colitis pre-clinical models of IBD, demonstrating improved survival and reduced colon inflammation and damage with reduced macrophage invasion and improved marker levels for thrombosis, uPAR and complement MAC when given as an acute prophylaxis in higher dose DSS colitis model and with acute exacerbations in a chronic lower dose DSS colitis model. Serpins are inhibitors that target active proteases with the potential to selectively target tissue sites of activated proteases (17,18). We have detected selective changes in inflammatory macrophage response, coagulation and inflammatory markers in the colon and submucosal vasculature. The inhibitory function of PEGSerp-1 is postulated to target tissue sites undergoing inflammation-induced protease activation, a *potential tissue targeting action*.

One distinct advantage of the serpin inhibitory mechanism, relative to non-covalent competitive inhibitors, is that once a productive encounter between the serpin and a protease has occurred, the inhibitory, covalent complex does not dissociate. Thus, durable inhibition is not predicated on local inhibitor concentration, which affects only the rate of engagement with a protease population. Many serpins have cofactors or ligands that can influence activity or localization. The results we have obtained suggest further tissue-specific targeting mechanisms may contribute to the localized effects of PEGSerp-1 that we have observed within the colonic tissue.

As DSS-induced colitis progresses, the levels of various innate and adaptive immune cells in the colonic tissue varies (2–8). In the acute prophylaxis 5% DSS and also in the acute and chronic delayed 2% DSS models, inflammatory cell infiltrates were significantly reduced by PEGSerp-1 treatment, closely paralleling significant decreases in inflammatory M1 macrophage cell invasion and crypt damage. Arg1 M2 anti-inflammatory macrophage were not modified by PEGSerp-1 treatment, suggesting a predominate M1 macrophage inhibition, without an increase in M2 macrophage activity, albeit providing a relative increase in M2 to M1 response through M1 suppression. Neutrophils and T cells are also reported to drive colitis, but in these models, there was a much smaller response to PEGSerp-1 with the exception of the chronic delayed, acute exacerbation 2% DSS mouse model, where PEGSerp-1 treatment significantly reduced CD4+T cells. These changes in the detected M1 counts in inflamed colon were paralleled in the submucosal vasculature in the 5% DSS acute prophylaxis model, but not in the chronic delay 2% DSS model.

The uPA protease and its uPA Receptor (uPAR) have both been linked to aggressive DSS colitis as well as associated with human biopsy specimens. Wild type Serp-1 protein has been previously demonstrated to bind and inhibit uPA and uPAR on human macrophages *in vitro* (32). uPA and uPAR activate plasmin which in turn activates matrix metalloproteinases that break down arterial collagen and elastin allowing macrophage to infiltrate the arterial wall. Serp-1 binding to the uPA/uPAR complex reduces macrophage invasion in inflammation mediated disease models, ranging from lupus lung hemorrhage and allograft transplants to SARS-CoV-2 and gamma herpes vasculitis and lung damage (33,34,42). The role of the receptor in this process is highlighted by the observation that in transplant models uPAR deficient allograft transplants were no longer protected by Serp-1 treatment in mouse aortic allografts (38). Similarly in a wound healing model Serp-1 treatment improved wound healing with reduced macrophage infiltrates, but with simultaneous uPAR antibody application, Serp-1 benefit was lost (38). Serp-1 mediated anti-inflammatory activity is thus dependent in part on blockade of the uPA/uPAR complex in other mouse models of vascular and wound healing. Upon treatment with PEGSerp-1, IHC showed uPAR to be reduced in the colon in the 4% DSS acute prophylaxis model and in the chronic delayed 2% DSS model, but not in the submucosal vessels in the colon samples from the acute 5% DSS prophylaxis model nor in the chronic delayed 2% DSS models. This is further supportive of a reduced infiltration of the colonic by inflammatory cells.

Activated serine proteases in the thrombotic cascade also drive microvascular thrombosis, micro-clots inducing ischemia, and complement activation that can cause further colon inflammation and damage. It was noted in prior vascular injury studies that Serp-1 treatment reduced arterial inflammation, and also reduced myocardial damage after coronary stent implant in a Phase 2A clinical trial (35). In the colitis studies reported here, PEGSerp-1 reduced colon and small vessel inflammatory cell responses in the colon wall with a reduction in fibrinogen, fXa and C5b/9 complement (MAC) detected by IHC. These broad effects of PEGSerp-1 suggest a targeted inhibition of vascular inflammation at sites of protease activation and organ damage that may be specific to the disease process. In prior work with a SARS-CoV-2 mouse infection model, uPAR appeared to have more specific response to PEGSerp-1 treatments (34) than seen here in the colitis models where a greater effect was detected in fibrinogen and fXa responses.

Mammalian serpins can represent up to 10% of circulating proteins and control thrombotic, thrombolytic and complement pathways central to normal physiologic functions in the mammalian body. Treatment with the mammalian serpin C1esterase inhibitor (C1Inh, SERPING1) reduced complement activation and is associated with reduced inflammation in colitis (15), whereas plasminogen activator inhibitor-1 (PAI-1, SERPINE1) blocks thrombolytic protease tissue type plasminogen activator (tPA) and increases inflammation in colitis (26). Mice deficient in PAI-1 and/ or mice treated with small molecule inhibitor of PAI-1 have reduced inflammatory colitis. Alpha 1 antitrypsin (AAT, SERPINA1) inhibits neutrophil elastase and inflammation, but is also reported to inhibit thrombotic proteases and treatment with AAT reduces colitis in mouse models (27,28). Antithrombin III (ATIII, SERPINC1) inhibits the thrombotic proteases and reduces inflammation in colitis (41). PEGSerp-1 as noted binds and inhibits tPA, uPA, plasmin fXa and thrombin as well as complement proteases (23–25,32–34), sharing many targets with mammalian serpins that have demonstrated benefit in reducing colitis. PEGSerp-1 is therefore postulated to provide anti-inflammatory activity in colitis predominately via blockade of fibrinogen, fXa, C5b/9 and potentially the uPA/uPAR, providing a potential targeted anti-thrombotic and anti-inflammatory activity in inflamed colon blood vessels.

Both acute aggressive 4% and 5% DSS colitis as well as lower grade more chronic 2%DSS colitis models were assessed. Treatment was assessed both as early prophylaxis as well as delayed treatment. In each case, early PEGSerp-1 treatment was most effective, demonstrating benefit in the acute prophylaxis as well as the chronic delayed 2%DSS model where an acute exacerbation in chronic colitis was mimicked by repeat DSS administration. Although no single animal model captures the complexity of IBD, oral administration of DSS is a widely used method for the chemical induction of colitis in mice. By varying the DSS concentration and duration of exposure, various models of colitis, including acute, chronic or relapsing were produced (36, 37). DSS induces colitis by inflicting damage on colon epithelial cells, thus compromising the barrier function of the epithelium. Disruption of the epithelial barrier results in the entry of luminal contents, including bacteria, into the deeper layers of the intestinal wall triggering an inflammatory immune response. The 2% DSS chronic delayed treatment model included in this project was designed to simulate the typical course of IBD, which is characterized by chronic remitting and relapsing inflammation of the gastrointestinal tract. Inflammation in the colon was established by administering 2% DSS for 5 days. The DSS was then removed to allow for a five-day recovery period before the DSS was reintroduced for 4 days to produce an acute flare of the disease, more closely simulating the clinical treatment of IBD than acute prophylaxis, as one will often see a delay between diagnosis and initiating treatments in severe colitis. 2% DSS was also administered for 7 days, with treatment beginning following the removal of the DSS in the acute delayed treatment model. In the acute delayed treatment model PEG Serp-1 did have some efficacy but less extensive when assessed by all markers. PEGSerp-1 treatment significantly reduced the extent of colon shortening, the depth of inflammatory cell infiltration and the number of CD3 positive T cells in the mucosa in each model. This study was limited to the use of male mice in a study designed to induce a more severe colitis assessing PEGSerp-1 treatment under more stringent conditions. With these studies PEGSerp-1 was given systemically at doses with previously proven efficacy. Given the beneficial results seen in this study, it will be of great interest to assess PEGSerp-1 treatment in further work in both male and female mice and with longer term more chronic colitis and at a range of doses.

When DSS-induced colitis progresses to a chronic state, key markers of disease severity in the acute stage, such as weight loss and diarrhea, no longer align with the severity of inflammation. Spleen size and colon length were assessed to evaluate the extent of inflammation. PEGSerp-1 treatment significantly reduced the extent of colon shortening, the depth of inflammatory cell infiltration and the number of CD3 positive T cells in the mucosa in each model. Mice in the PEGSerp-1 group generally had smaller spleens by visual inspection but not significant by quantification. There was also a marked reduction in markers for thrombosis as well as complement in the colon and the submucosal vessels. uPAR was reduced in the colon but less marked reduction was seen in the colon vasculature suggesting a greater effect on the thrombotic and complement cascades than uPAR in colitis and associated vasculitis. Further analyses of effects on coagulation parameters will be of interest in future work.

RT-PCR analysis of gene expression in colon and heart samples, from the chronic delayed 2% DSS model, detected altered uPAR expression in the colon with a strong correlation with colon damage (Figure 9, Panel 2A) and confirmed DSS induced upregulation of inflammatory response pathways, specifically fXa (Figure 9, Panel 2B) and C3 complement (Figure 9, Panel 2C), within the tissue. Effects of PEGSerp-1 treatment on gene expression in the 2% chronic delayed model detected a significant increase for PAI-1 in the colon (Figure 9, Panel 2E), and a trend for increased gene expression for uPAR (Figure 9, Panel 2D) and C1Inh (Figure 9, Panel 2F). There were otherwise non-significant directional changes in gene expression as observed in the IHC analysis on histopathological analysis. Overall, there were similar trends in gene expression and IHC analyses as has been seen in prior work with the SARS-CoV-2 mouse models where localized significant changes in coagulation and complement were also detected again supporting tissue targeted serpin treatment. One technical factor that may have impacted the present study was a delayed analysis of gene expression. In the 2% chronic delayed colitis model, gene expression was measured at 20 days, 10 days after reintroduction of 2% DSS to the drinking water and 7 days after the final delayed dose of PEGSerp-1 which was initiated on day 10 when DSS was re-introduced. The RNA profile may thus not be representative of peak responses to PEGSerp-1 treatments. In addition, the colon contains multiple differing cells as evidenced by the associated colon vascular and lymphatic tissues in the colon which may also vary in their responses to DSS treatment and PEGSerp-1 treatments.

In this project, murine DSS-induced colitis was used to evaluate systemic PEGSerp-1 as a treatment for IBD, colon inflammation and damage and associated vascular inflammation and evidence for thrombosis. Three studies were conducted representing three different disease and treatment models. When administered daily via intraperitoneal injection as either a preventative or delayed treatment, PEGSerp-1 improved outcomes in mice when given early as an acute prophylaxis in 4% and 5% DSS models and in a model of 2% DSS colitis with acute recurrence. PEGSerp-1 was less effective when given 7 days after inducing colitis with 2% DSS. Efficacy of PEGSerp-1 treatment was closely associated with reduced colon damage and inflammation as well as reduced macrophage invasion. The doses used for PEGSerp-1, at 2µg per mouse per day, have proven effective in lupus lung hemorrhage and SARS-CoV-2 mouse models and are, in general, approximately 100-1000 fold lower than doses used for prior mammalian serpin treatments. Further analysis will determine the efficacy of PEGSerp-1 at a wider range of doses and duration of treatments for acute exacerbations and with comparison analysis for current approaches to treatment.

**We report here investigation of the use of a purified serpin, originally of viral origin, given early to treat excess systemic inflammation and coagulation pathway activation in a mouse model of IBD. This improves survival and reduces tissue damage, supporting further investigation of PEGSerp-1 as a tissue-targeting treatment for IBD.**

## Materials and Methods

### 2.2 PEGSerp-1: Protein purification and PEGylation

Serp-1 (m008.1L; NCBI Gene ID# 932146) was expressed in a Chinese hamster ovary (CHO) cell line in (33–35,42) as previously described to GMP-compliant standards. This was >95% pure, as determined by Coomassie stained SDS-PAGE and reverse-phase HPLC and endotoxin-free by limulus amebocyte lysate assay. For PEGylation, Serp-1 was incubated with mPEG-NHS (5 K) (Nanocs Inc., #PG1-SC-5k-1, NY) in PBS buffer (pH 7.8) at 4^◦^C overnight according to standard PEGylation protocols (33,34). PEGSerp-1 was purified by FPLC using an ÄKTA pure protein purification system (Cytiva) using a Superdex-200 column and found to have a mean number of 5 PEGylation sites (m5)(33,34). PEGSerp-1 demonstrated binding to active uPA as assessed on Western blot (Supplemental Figure S3 and Supplemental Table S1).

### 2.3 Colitis induction and treatment

All animal protocols were reviewed and approved by the Institutional Animal Care and Use Committee (IACUC) of Arizona State University (# 20-1761R). Dextran sulfate sodium (DSS) is a sulfated polysaccharide which can be delivered orally to induce experimental colitis in mice (36,37). Disease severity and course can be modified by varying DSS concentration from 1.5 to 5%, with varying duration of DSS exposure (36,37). C57BL/6 mouse strains develop a chronic colitis on exposure to DSS. Male mice have a more aggressive DSS induced colitis and were used for these initial studies. C57BL/6J mice used in this study were bred at the Biodesign Institute by Dr. Wei Kong’s team.

Colitis was induced when the mice reached 7-8 weeks of age by supplementing the drinking water with DSS (molecular weight, 36-50 kD; MP Biomedicals, Solon, OH). The DSS concentration, duration of DSS exposure and timing of treatment initiation were varied to produce three experimental models (Figure 1, Table 1) based upon DSS percentage and treatment, *eg.* 5% for 6 days and 4% DSS for 5 days with acute prophylaxis treatments with PEGSerp-1 starting on the day DSS was introduced in the drinking water and continued for 14 days (Figure 1, Group A), 2% DSS for 7 days with acute delayed PEGSerp-1 given starting 7 days after adding DSS to the water and continued for 7 days (Figure 1, Group B) and finally an initial 5 days 2% DSS followed by a 5 days rest and repeat 2% DSS addition to the drinking water starting on day 10, with PEG Serp-1 treatment starting on day 10 when 2% DSS is reintroduced to the drinking water and continued for 4 days (Figure 1, Group B). Treatments were given by daily intraperitoneal (IP) injections of either 100 μL saline or 2µg of PEGSerp-1 (100 ng/mg bodyweight) in 100 μL of saline. Across all studies, mice were randomly assigned to treatment groups before DSS induction (Table 1).

**Table 1.** Summary of experimental C57BL/6J mice included in this study.

|  | DSS<br>Concentration | DSS<br>Exposure | Treatment<br>Duration | Treatment | Number<br>of mice | Total study<br>duration |
| --- | --- | --- | --- | --- | --- | --- |
| healthy control | No DSS |  |  |  | 4 | 14 days |
| Acute Prophylaxis | 5% | Days 0-6 | Day 0-13 | Saline | 5 | 14 days |
|  |  |  |  | PEGSerp-1 | 5 |  |
|  | 4% | Days 0-5 | Day 0-13 | Saline | 5 | 14 days |
|  |  |  |  | PEGSerp-1 | 5 |  |
| Acute Delayed | 2% | Days 0-7 | Days 7-13 | Saline | 10 | 14 days |
|  |  |  |  | PEGSerp-1 | 10 |  |
| Chronic delayed | 2% | Days 0-5<br>and<br>10-13 | Days 10-19 | Saline<br>PEGSerp1- | 10,<br>10 | 20 days |

Two initial studies were conducted where mice were administered 5% or 4% w/v DSS and prophylaxis with PEGSerp-1 tested for efficacy. The DSS concentration was reduced to 2% weight/volume (w/v) in subsequent studies to simulate chronic colitis with lower mortality. Acute delayed treatment and chronic delayed treatments were assessed in the 2% w/v DSS model with treatments starting at 7 days (Figure 1B) or at 10 days when 2% DSS was re-introduced into the drinking water. Within the chronic delayed treatment model, 3 mice were removed from each treatment group due to severe disease (defined by 25% body weight loss) prior to the beginning of treatment on day 10. Four control mice, without DSS exposure were examined. Mice were weighed daily and euthanized if any signs of distress.

### 2.4 Tissue collection and Histological Analysis

At the time of tissue collection, the colon and cecum were removed and the length recorded as shown in Figure 3. Spleens were examined after excision and were measured in the chronic delayed treatment model. Collected tissues were fixed in 10% neutral-buffered formalin, processed and then embedded in paraffin. Colon tissue was cut into 5 μm thick sections for immunohistochemical (IHC) staining (33,34,36,37).

The extent of epithelium /colon damage was evaluated at 2x magnification by dividing the length of tissue with damaged architecture (depletion of or loss of crypt structure) by the total length of the section measured (Figure 4A-B). The degree of inflammatory cell infiltration was also assessed at higher magnification (20x magnification) by dividing the depth of cell infiltrate from the submucosa and dividing by the total thickness of the section (Figure 2F-I). Crypt architecture was measured at 2X by dividing the depth of the longest crypt in a colitis specimen by the total colon wall thickness.

### 2.5. Immunohistochemical staining (IHC)

Immunohistochemical (IHC) staining analysis was performed to assess immune cell responses as well as inflammatory thrombotic and thrombolytic pathway markers. Positively stained cells were counted at 20x magnification in 3 high power fields (HPF) for each set of mouse colon samples (*e.g.* each individual mouse). Immune cell markers assessed included iNOS+ M1 and Arginase 1+ M2 for macrophage; CD3 and CD4 for T lymphocytes, and Ly6G for neutrophils: nonspecific T lymphocytes (CD3, Abcam ab5690, 1:100 and CD4, Abcam, ab183685, 1:1,000), M1 macrophages (iNOS, Abcam ab15323, 1: 200), M2 macrophage arginase-1 (Cell Signaling, 93668, 1:200 and neutrophils (Ly6G, Invitrogen 1960181, 1:100). For inflammatory and coagulation markers, fibrinogen antibody (XXX, 1:200), factor X antibody (Abcam # ab34269, 1:200), and uPAR (R&D Systems, AF534,1:100) as well as complement MAC (C5b/9, Abcam, ab 55811, 1:200). HRP conjugated secondary antibodies used were Anti-rabbit IgG (Jackson 111-035045) and goat anti-rat IgG (Jackson 112-035-143) at a dilution of 1:500 Cat: A110-109P along with ImmPACT DAB (Vector Labs, USA) to visualize the antigens. HRP-conjugated secondary antibody given alone without primary antibody was used as negative control for each stain. Antigens were revealed with ImmPACT DAB (Vector Labs, USA), counterstained with Gil’s formula #3 Hematoxylin and mounted with Cytoseal XYL. Sections were examined using an Olympus BX51 microscope with 2x–100x objectives, a Prior ProScan II stage and Olympus DP74 CMOS camera and cellSens software analysis system.

An independent, blind pathology score was also performed (SG), with score based on colon inflammatory cell invasion and loss of crypt architecture with apoptosis (43). If the colon tissues were clean without significant inflammation or edema, the Score assigned was 0+. For colon with mild to moderate inflammation and damage, Score assigned was 1-2+. For sections with extensive colon inflammation and loss of architecture, Score assigned 3+.

### RNA isolation and qPCR

Quantitative polymerase chain reaction (qPCR) was performed to determine if the expression levels of cytokines, proteases and serpins were modified by PEGSerp-1 treatment (27). Samples were assayed for gene expression in colon and heart tissues isolated from the chronic delayed 2% DSS model (Figure 1, Group C), with and without PEGSerp-1 treatment as well as in normal heart isolates. Colon and heart samples from controls or 2% DSS colitis model mice were collected 20 days after reintroduction of 2% DSS and frozen. Samples were also collected from formalin fixed, paraffin embedded (FFPE) colon samples in the 5% DSS acute prophylaxis model but yields were low and data was not considered reliable.

For RNA extraction, tissues were harvested in RNA STAT-60 (Tel-Test) and disrupted by bead homogenization (Omni, INC). Isolated mouse lungs were transferred to Trizol (Thermo Fisher Scientific). RNA was extracted using the Qiagen RNAeasy Mini Kit (Qiagen, 74106), for frozen tissues and using the Qiagen RNAeasy kit (Qiagen 73504) for FFPE specimens, according to the manufacturer’s instructions. Briefly, 0.2 mL chloroform was added to 1 mL of RNA STAT-60, samples were mixed for 30 seconds, kept on ice for 5 min and centrifuged 12,000g for 15 minutes at 4 °C. 0.5 mL of 100% ethanol was added to aqueous phase (RNA), samples were kept for 5 min at 4 °C and centrifuged as above. The final RNA pellet was resuspended in molecular biology grade water (Sigma). Total RNA was reverse transcribed with an oligo (dT) primer and a Moloney murine leukemia virus reverse transcriptase (MMLV RT, TakaraBio). cDNA was analyzed by quantitative PCR amplification using SYBR Green qPCR Master Mix (QuantaBio) on a Bio-Rad CFX96 Real-Time PCR Detection System. Primers were designed to amplify mRNA-specific sequences, and analysis of the melt-curve confirmed the amplification of single products. Uninfected samples were used as controls. Relative expression was normalized to GAPDH. Primer sequences used are provided in Supplemental Table S2. Primers were designed using the IDT website, PrimerQuest Tool, for qPCR and confirmed in the literature.

Primers used for qPCR included Gapdh, Fx, Tpa, Upa, Upar, Pai-1, Il-1, Il-6, Vegf, C3, C5, C1inh (Supplemental Table S2). Real time qPCR was analyzed on a BioradCFX96 Real Time PCR array with C1000 Touch Thermal cycler.

### 2.5 Statistical analyses

Three colon sections were analyzed for each mouse and mean values of measured changes for IHC analysis were used all statistical analyses. Data were analyzed using StatView (SAS Institute, Inc.) or GraphPad Prism (GraphPad Software, San Diego, CA). Kaplan-Meier cumulative Survival analysis was performed. Differences in colon length (gross analysis and histology at 2X) and altered histology and IHC analyses (20X) between multiple groups were evaluated by analysis of variance (ANOVA) with Fisher’s least significant difference (LSD) post hoc analysis between individual subgroups. P < 0.05 was considered significant. Changes between two treatment groups were analyzed by unpaired Student’s t tests.

## Supplemental Figures

**Supplemental Figure S1.**
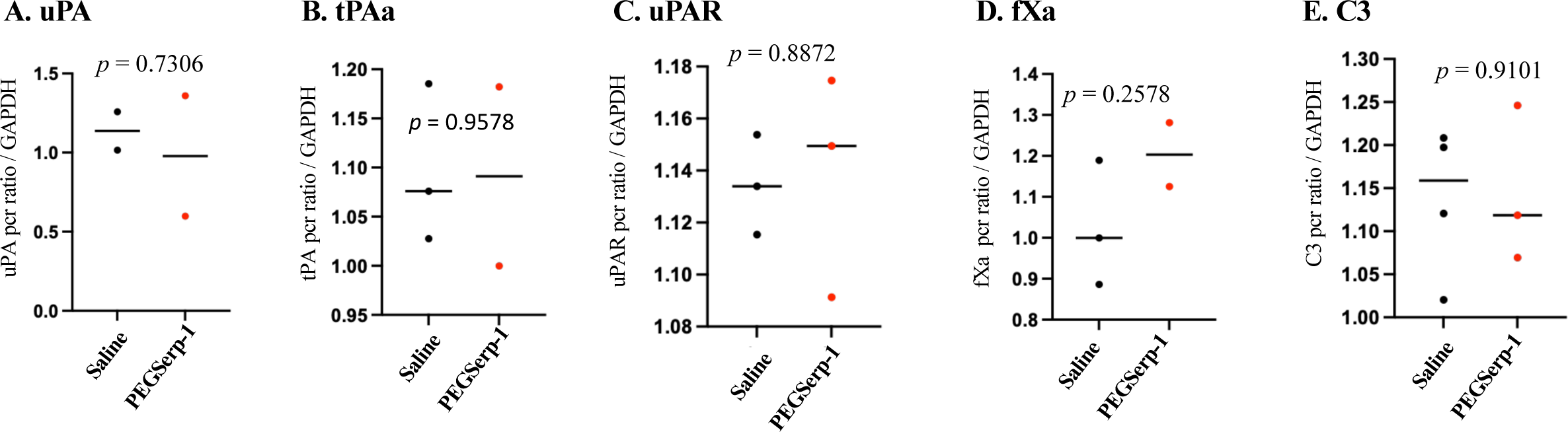
RT-PCR of Colitis tissue - 2% DSS Chronic delayed

**Supplemental Figure S2.**
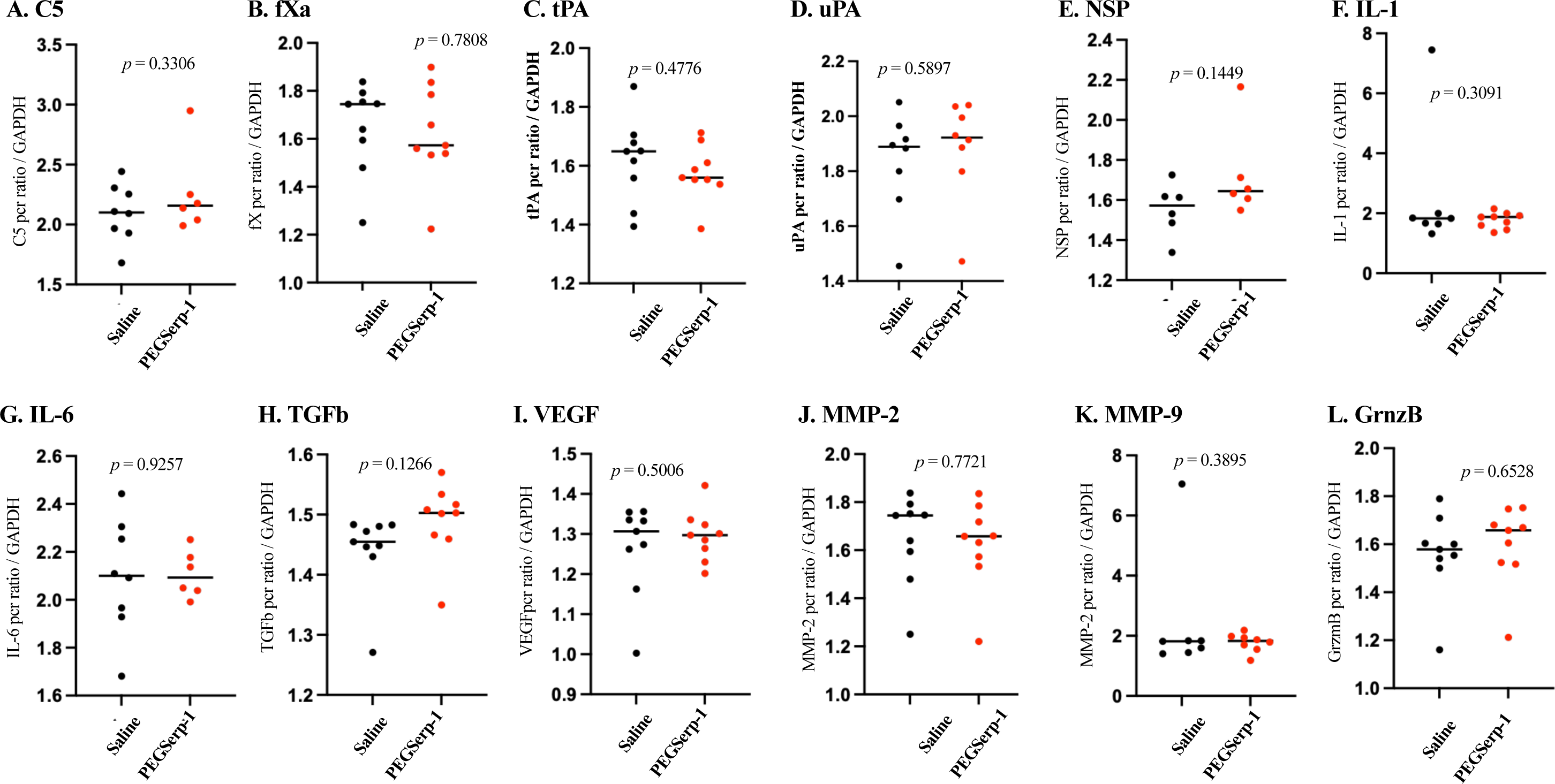
RT-PCR analyses of heart tissues – 2% Chronic delayed

**Supplemental Figure S3.**
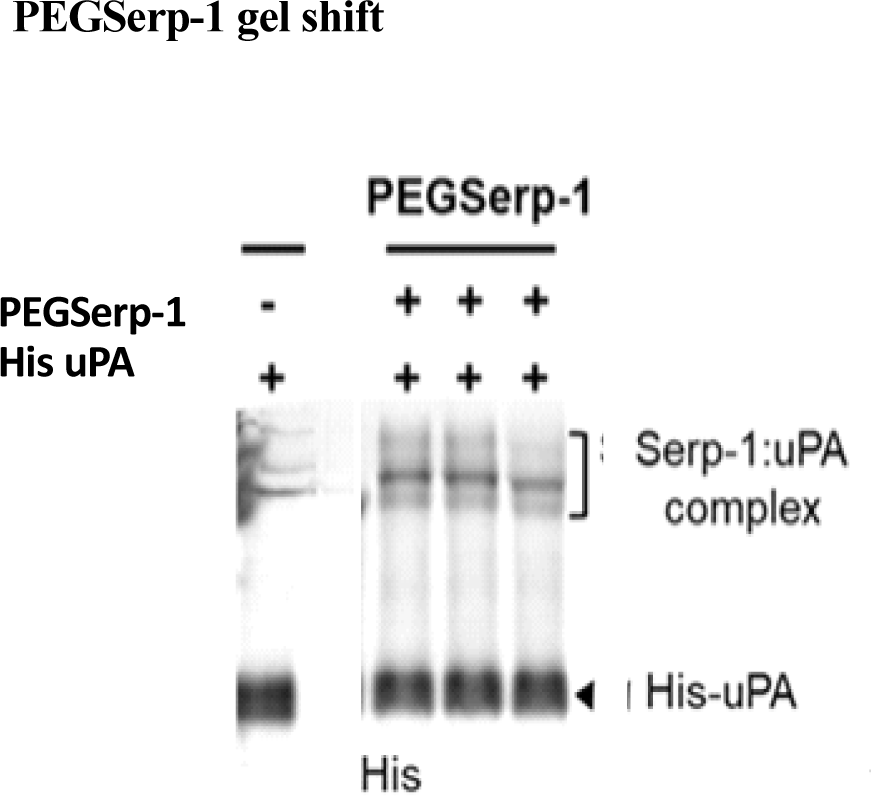
PEGylation and purification of PEGSerp-1

**Supplemental Table S1.** Mass Spectrometry analysis of PEGSerp-1 binding in lung hemorrhage model.

| Accession | Description | Symbol | # Peptides |
| --- | --- | --- | --- |
| P01027 | Complement C3 | C3 PE | 34 |
| B7FAV1 | Filamin, alpha | Flna PE | 23 |
| P60710 | Actin, cytoplasmic 1 | Actb PE | 17 |
| E9PV24 | Fibrinogen alpha chain | Fga PE | 12 |
| P20918 | Plasminogen | Plg PE | 11 |
| P01029 | Complement C4-B | C4b PE | 9 |
| P98086 | Complement C1q subunit A | C1qa PE | 4 |
| P04186 | Complement factor B | Cfb PE | 4 |
| Q8CG14 | Complement C1s-A | C1sa PE | 4 |
| P14106 | Complement C1q subunit B | C1qb PE | 3 |
| Q8CG16 | Complement C1r-A | C1ra PE | 3 |
| Q02105 | Complement C1q subunit C | C1qc PE | 2 |

\*Average Xcorr >4.0

**Supplemental Table S2.**
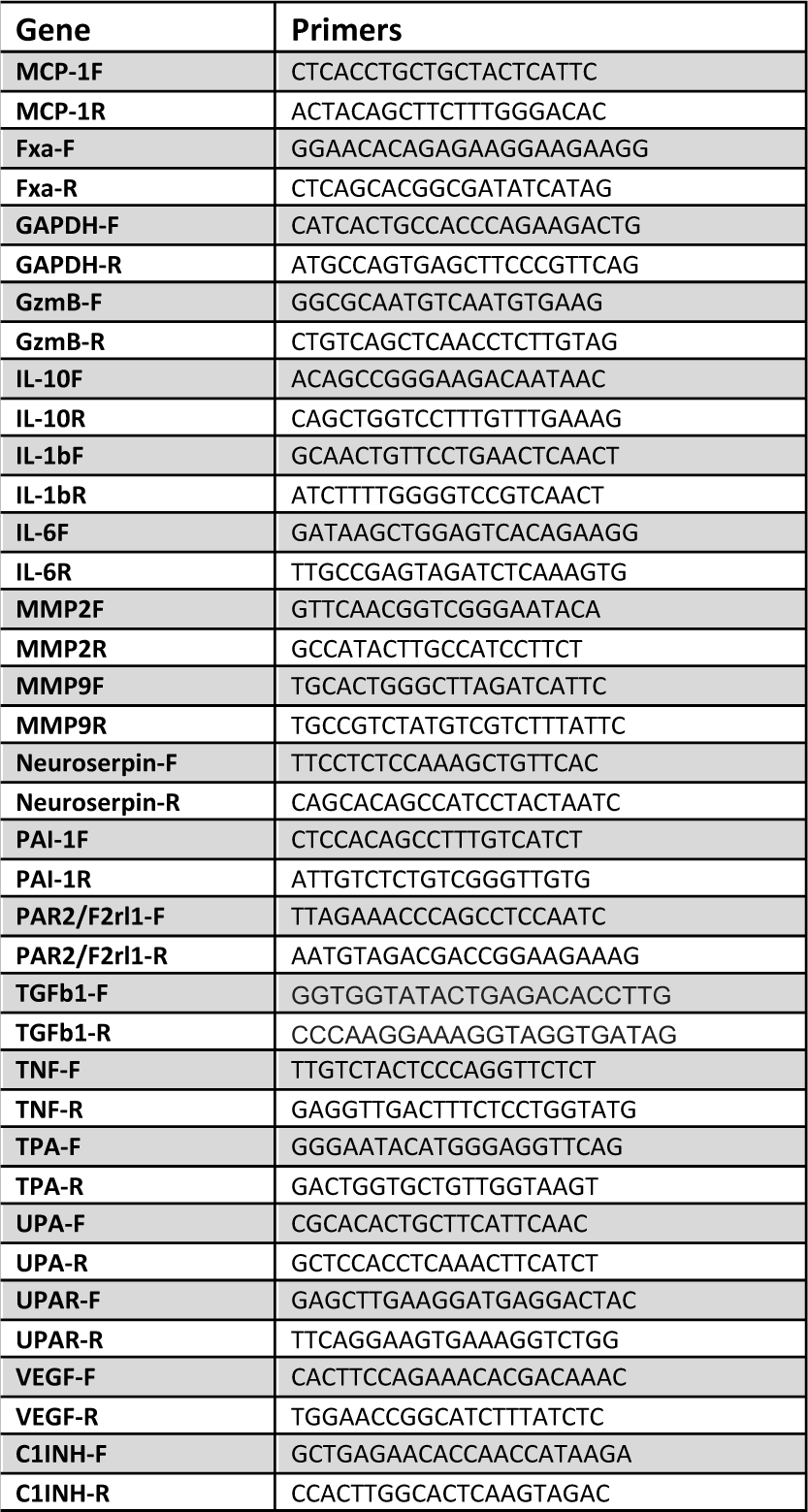
primers per EMM paper.

